# Astrocyte GluN2C NMDA receptors control basal synaptic strengths of hippocampal CA1 pyramidal neurons in the *stratum radiatum*

**DOI:** 10.1101/2021.05.28.446253

**Authors:** Peter H. Chipman, Alejandra Pazo Fernandez, Chi Chung Alan Fung, Angelo Tedoldi, Atsushi Kawai, Sunita Ghimire Gautam, Mizuki Kurosawa, Manabu Abe, Kenji Sakimura, Tomoki Fukai, Yukiko Goda

## Abstract

Experience-dependent plasticity is a key feature of brain synapses for which neuronal N-Methyl-D-Aspartate receptors (NMDARs) play a major role, from developmental circuit refinement to learning and memory. Astrocytes also express NMDARs although their exact function has remained controversial. Here we identify a circuit function for GluN2C NMDAR, a subtype highly expressed in astrocytes, in layer-specific tuning of synaptic strengths in mouse hippocampal CA1 pyramidal neurons. Interfering with astrocyte NMDAR or GluN2C NMDAR activity reduces the range of presynaptic strength distribution specifically in the *stratum radiatum* inputs without an appreciable change in the mean presynaptic strength. Mathematical modeling shows that narrowing of the width of presynaptic release probability distribution compromises the expression of long-term synaptic plasticity. Our findings suggest a novel feedback signaling system that uses astrocyte GluN2C NMDARs to adjust basal synaptic weight distribution of Schaffer collateral inputs, which in turn impacts computations performed by the CA1 pyramidal neuron.

## Introduction

Plasticity is a fundamental feature of neuronal connections in the brain, where experience-dependent changes in synaptic strengths over different time scales are crucial for a variety of processes ranging from neural circuit development, circuit computations to learning and memory (Feldman and Brecht, 2005; Bliss and Collingridge, 1993; Abbott and Regehr, 2004; Collingridge et al., 2010; Nicoll, 2017). Deciphering how neurons dynamically express different forms of synaptic plasticity while ensuring the optimal performance of the circuit remains a key challenge (Vitureira and Goda, 2013; Turrigiano, 2017; Nicoll, 2017; Brunel, 2016). In particular, the distribution of synaptic strengths is thought to reflect the capacity or the state of information storage of neural circuits (Barbour et al., 2007; Buzsáki and Mizuseki, 2014; Bromer et al., 2018). A better understanding of the cellular mechanisms that regulate synaptic strength distribution could therefore provide novel insights into the basis for the effective execution of circuit operations (Barbour et al., 2007).

The N-methyl-D-aspartate receptors (NMDARs) are a major ionotropic glutamate receptor type mediating excitatory synaptic transmission (Paoletti et al., 2013; Sanz-Clemente et al., 2013; Hansen et al., 2018). NMDARs are heteromeric assemblies consisting of four subunits, and seven NMDAR subunit genes have been identified that fall into three subfamilies: *Grin1* encoding the obligatory GluN1 subunit, four *Grin2* genes encoding GluN2A-D, and two *Grin3* genes encoding GluN3A-B (Paoletti et al., 2013; Hansen et al., 2018). The differing subunit compositions confer NMDARs with distinct biophysical and pharmacological properties and contribute to their diverse biological activities. NMDARs are expressed throughout the central nervous system (CNS) and are crucial for normal brain function and plasticity. In particular, NMDARs that are typically composed of two GluN1 and two GluN2 subunits and are present postsynaptically have been extensively studied for their role in memory mechanisms (Paoletti et al., 2013; Sanz-Clemente et al., 2013; Nicoll 2017). NMDARs are also involved in pathological conditions such as epilepsy, ischemia, and traumatic brain injury (Hansen et al., 2017). In stark contrast to the multifaceted functions of neuronal NMDARs, very little is known of NMDAR functions in glial cells in the CNS. Amongst the better characterized glial NMDARs, NMDARs in oligodendrocytes play a role in axonal energy metabolism (Saab et al., 2016) and exacerbate pathological conditions (Karadottir et al., 2005). In astrocytes, NMDARs have been long thought to be absent except following ischemia *in vivo* or under anoxia *in vitro* that promote their aberrant expression (Krebs et al., 2003; Gottlieb and Matute, 1997). Nevertheless, there are reports of GluN2C mRNA expression in astrocytes in the adult brain (Watanabe et al., 1993; Karavanova et al., 2007; Ravikrishnan et al., 2018), and several studies provide physiological evidence in support of astrocyte NMDAR expression under non-pathological conditions (Schipke et al., 2001; Serrano et al., 2008; Lalo et al., 2006; Letellier et al., 2016). The precise function for astrocyte NMDARs, however, has remained a matter of debate (e.g. Kirchhoff, 2017), and the role for astrocyte GluN2C has not yet been identified.

Astrocytes regulate a diverse set of essential brain activities, from neurovascular coupling, metabolic support, and ionic homeostasis (e.g. Attwell et al., 2010; Giaume et al., 2010; Simard and Nedergaard, 2004) to synaptic connectivity and function (e.g. Araque et al., 2014; Clarke and Barres, 2013). Recent studies have highlighted a role for astrocytes in the bidirectional control of basal synaptic transmission (Panatier et al., 2011; Perea and Araque; 2007; DiCastro et al., 2011; Martin-Fernandez et al., 2017; Schwarz et al., 2017), which likely involves signaling via their perisynaptic processes (Bindocci et al., 2017; Bazargani and Attwell, 2016). Moreover, mounting evidence suggests that astrocytes are not only capable of either simply potentiating (Panatier et al., 2011; Perea and Araque, 2007; Jourdain et al., 2007) or depressing synaptic transmission (Martín et al., 2015; Pascual et al., 2005), but a single astrocyte, which contacts tens of thousands of synapses, can concurrently mediate bi-directional synapse regulation at separate synaptic contact sites (Schwarz et al., 2017; Covelo and Araque, 2018). Astrocytes likely regulate neural circuit functions at multiple levels, as suggested by observations of both local and global astrocyte signaling that are triggered in a synapse, neuron, or circuit-specific manner (Martín-Fernandez et al., 2017; Martín et al., 2015; Chai et al., 2017; Deemyad et al., 2018; Dallérac et al., 2018; Santello et al., 2019). However, despite these advances, the basic mechanisms by which astrocytes detect and adjust synaptic transmission and the extent of their impact on local synaptic circuit activity are not fully understood.

Astrocytes express a plethora of neurotransmitter receptors and membrane transporters that are thought to modulate synaptic transmission (Araque et al., 2014; Bazargani and Attwell, 2016), including perisynaptic astrocyte glutamate transporters that influence the magnitude and kinetics of postsynaptic glutamate receptor activation and membrane depolarization (Pannasch et al., 2014; Murphy-Royal et al, 2015) and astrocyte metabotropic glutamate receptors (mGluRs) whose activation promote gliotransmitter release to provide feedback control of synaptic transmission (Panatier et al., 2011; Schwarz et al., 2017; Covelo and Araque, 2018; Araque et al., 2014; Bazargani and Attwell, 2016). Given the often shared expression of neurotransmitter receptors and transporters between astrocytes and neurons, an important challenge is to decipher the synaptic activity-dependent functions of astrocyte receptors and transporters independently of the functions of their neuronal counterparts. Notably, many of the prior studies have focused on pathological brain states (Krebs et al., 2003; Gottlieb and Matute, 1997) or used young brain tissue (e.g. Panatier et al., 2011) or culture preparations (e.g. Schwarz et al., 2017) in which the expression pattern of the astrocyte receptors and channels differ substantially from the adult brain in basal conditions (Sun et al., 2013; also see Boisvert et al., 2018). Consequently, the cellular basis and the network consequences of astrocyte-synapse interactions mediated by the astrocyte receptors and channels in the healthy adult brain remain to be clarified.

In a previous study, we identified a role for astrocyte signaling in regulating synaptic strength heterogeneity in hippocampal neurons, which involved astrocyte NMDARs (Letellier et al., 2016). Heterogeneity was assessed by comparing the presynaptic strengths of two independent inputs to pyramidal neurons, which was estimated by the paired-pulse ratio (PPR) of excitatory postsynaptic current (EPSC) amplitudes, a parameter inversely related to presynaptic release probability (Dobrunz and Stevens, 1997). Although basal PPRs were uncorrelated between the two inputs, surprisingly, inhibiting astrocyte NMDARs promoted the correlation of PPRs. This indicated that astrocyte NMDARs contributed to enhancing the differences in basal presynaptic strengths represented by the two different inputs (Letellier et al., 2016). Such pair-wise comparison of PPRs, however, is limited to providing an indirect measure of variability, and it remains unclear how astrocyte NMDARs control the overall shape of the PPR distribution. Furthermore, the crucial roles for neuronal NMDARs in regulating synaptic transmission and plasticity have confounded the interpretation of astrocyte NMDAR functions, which still remain enigmatic (Kirchhoff, 2017). Here, we further investigated astrocyte NMDAR-dependent regulation of synaptic strength in hippocampal CA1 pyramidal neurons in adult mice, focusing on the mode of regulation of presynaptic strength distribution, the NMDAR type responsible, and probing the potential layer specific regulation in area CA1. Our findings identify a role for astrocyte GluN2C NMDARs in maintaining broad presynaptic strength diversity specifically in the *stratum radiatum* (SR) by enhancing both strong and weak synapses. Mathematical modeling suggests that presynaptic strength diversity strongly influences the expression of long-term synaptic plasticity, which in turn, is important for network stability and learning and memory (Malenka and Bear, 2004; Collingridge et al., 2010; Royer and Paré, 2003; Zenke and Gerstner, 2017). Our findings suggest astrocyte GluN2C NMDARs as a key player in linking the regulation of synaptic strength distribution to the expression of synaptic plasticity that promotes optimal circuit performance.

## Results

### NMDAR-dependent regulation of PPR variability in hippocampal CA1 pyramidal neurons

In order to characterize the properties of astrocyte NMDAR-dependent modulation of synaptic strength, here we devised a simplified assay using a single stimulating electrode to monitor PPR variability of Schaffer collateral synapses. Hippocampal slices were prepared from adult mouse brain (P60-120), and EPSCs to a pairwise stimulation (50 ms inter-stimulus interval) of Schaffer collateral axons were recorded in CA1 pyramidal neurons (Figure 1A; Figure S1). MK-801 (1 mM) was infused into the postsynaptic cell via the patch pipette, and after 10-15 min but prior to starting the experiment, Schaffer collaterals were stimulated at 0.1 Hz for at least 45 times to pre-block synaptic NMDARs. This resulted in over 92% inhibition of synaptic NMDAR currents (Bender et al 2006) (Figure S1; see methods). Upon confirming the block of postsynaptic NMDARs, the baseline EPSC recordings were taken; subsequently NMDAR inhibitors were bath applied to test their effects on PPR variance. Under the block of postsynaptic NMDARs, any effects of NMDAR inhibitors, if observed, would be expected to reflect the influence of NMDARs present either in astrocytes or the presynaptic neurons. Notably, in mature brains which we used for our experiments, the contribution of presynaptic NMDARs in general had been suggested be minimal (Shih et al., 2013). Upon bath applying MK801 (50 µM) or AP5 (50 µM), the population variance (δ^2^) of PPRs was reduced without a significant change in the mean (x̅) (Figure 1B-D; MK801 δ^2^ p=.016, x̅ p=.856; AP5, δ^2^ p=.047; x̅ p=.162). This overall decrease in PPR variance, when examined at the level of individual inputs, was associated with a change in PPR (ΔPPR) that was negatively correlated to the baseline PPR: PPR of some inputs increased while others decreased by the NMDAR antagonist application (Figure 1E-G). Furthermore, consistent with a lack of change in the mean PPR upon blocking NMDARs, a linear fit to the data intercepted the x-axis near the baseline mean PPR (x-axis intercept: MK801 = 1.97; AP5 = 1.95).

**Figure 1.**
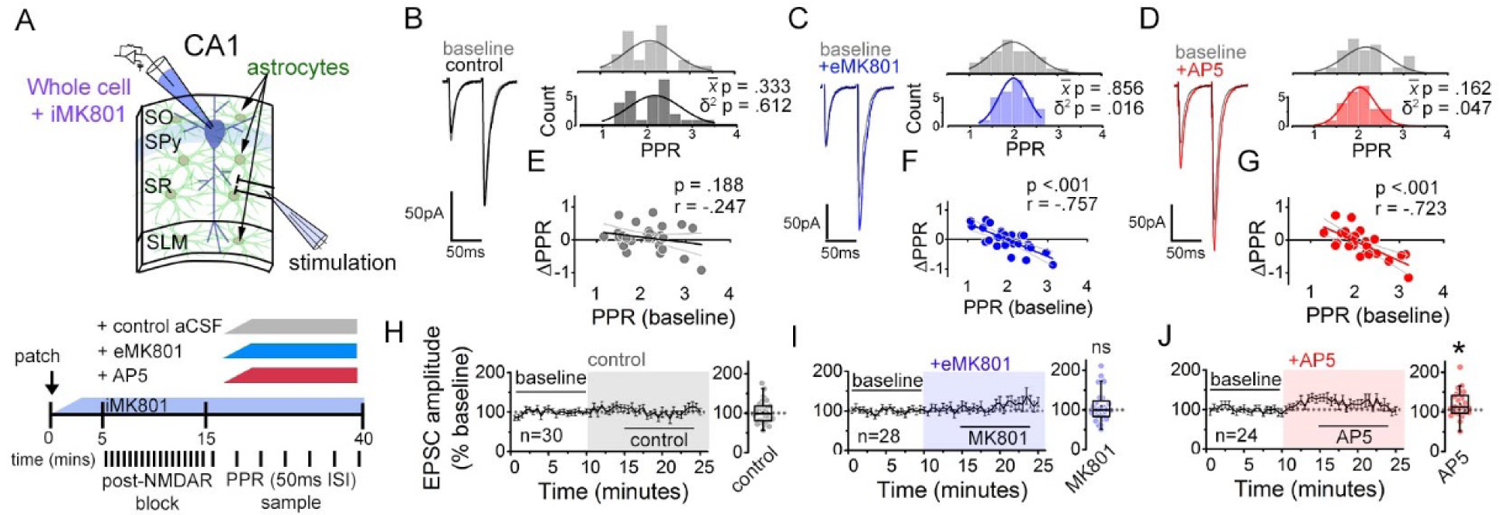
NMDARs contribute to synaptic strength diversity of Schaffer collateral synapses. (**A**) Experimental scheme to assess the role of non-postsynaptic NMDARs. MK801 was perfused through the patch pipette to the CA1 pyramidal neuron for 10-15 min, and pairs of stimuli (Δt = 50 ms) were applied at least 45 times at 0.1Hz to pre-block postsynaptic NMDARs prior to bath applying NMDAR antagonists (bottom). (**B-D**) Left: representative EPSC traces (average of 20 sweeps) to pairs of pulses (Δt = 50 ms) applied to Schaffer collateral axons before (baseline) and after wash-on of vehicle control (*B*), MK801 (50 µM) (*C*), or AP5 (50 µM) (*D*). Right: histograms of the PPRs recorded during the baseline and vehicle or drug application periods were fit with single Gaussian curves. The p values for the comparison of the baseline to the experimental periods for the population mean (x̅) and variance (δ^2^) were obtained by two-tailed paired sample t-test and one-tailed f-test for equal variances, respectively. (**E-G**) Scatterplots of the change in PPR vs. the initial PPR in vehicle control (*E*), MK801 (*F*), or AP5 (*G*) where ΔPPR = PPR_experimental_-PPR_baseline_; linear fits, Pearson’s correlation coefficients (r) and p values are as indicated. X intercept: MK801 = 1.97, AP5 = 1.95. (**H**-**J**) Plots of normalized EPSC amplitudes before and during the application of NMDAR antagonists (shaded area); n, number of inputs examined for each experiment. Baseline and experimental periods are indicated (black bars). Right: summary bar graph. * p<0.05, Mann-Whitney U-test. Control = 30 inputs, 15 cells, 13 mice; MK801 = 28 inputs, 14 cells, 9 mice; AP5 = 24 inputs, 12 cells, 11 mice.

We also tested for the contribution of GluN2B NMDAR and also of mGluR5 in maintaining PPR variance. Bath applying Ro25-6981 (10 µM), a GluN2B-specific NMDAR antagonist (Fischer et al., 1997), and MPEP (25 µM), an mGluR5-specific antagonist (Gasparini et al., 1999), had little effect on PPR variance (Figure S2A-C; Ro256981 p=.362, MPEP p=.349). MPEP, however, significantly increased the mean PPR (Figure S2C; p=.033), which is consistent with a reported role for mGluR5 in presynaptic strength regulation (Panatier et al 2011).

We next examined whether the changes in PPR elicited by the extracellularly applied NMDAR inhibitors accompanied changes in the EPSC amplitude and the coefficient of variation (CV) of EPSC amplitude, the latter as an additional measure of the change in presynaptic release probability (Malinow and Tsien, 1990; Larkman et al, 1992). On average, bath applied MK801 did not substantially alter the amplitude of the first EPSC to the pair of stimuli relative to control slices, despite a small tendency for an increase with a time delay (Figure 1I; p= .657). MK801 also had little effect on CV^-2^ of EPSC amplitudes (Figure S3B; p= .991), although the responses of individual inputs to MK801, albeit small in magnitude, were positively correlated to CV^-2^ (Figure S3D; Pearson’s r = .420, p = .026) and negatively correlated to PPR (Figure S3H; Pearson’s r = −.474, p = .011). These data are consistent with the possibility that the observed reduction in PPR diversity by MK801 application is associated with normalization of presynaptic release probability.

In contrast to MK801, AP5 significantly increased EPSC amplitudes within 10 min of application (Figure 1J; p = .026), and the increase in CV^-2^ (Figure S3B; AP5 p = .003) suggested that the change in EPSC amplitude involved a presynaptic change. Notably, Ro25-6981 that had no effect on PPR variance, did produce a significant increase in CV^-2^ (Figure S2H; CV^-2^ p = .002), and produced a small but non-significant increase in EPSC amplitudes (Figure S2E; EPSC p = .261), which was consistent with the observed increase in PPR (above). None of the NMDAR antagonists caused substantial changes in the amplitude or frequency of spontaneous EPSCs (sEPSCs) or evoked EPSC waveforms (Figure S4).

We next sought to confirm the NMDAR-dependent modulation of presynaptic strength diversity as described in our previous study by comparing PPRs across two independent Schafer collateral inputs that converge onto a target CA1 pyramidal neuron (Letellier et al., 2016), and to observe the time course of PPR population variance change by NMDAR antagonists. In this method, the absolute difference of PPRs between the two inputs (PPR disparity) is taken as a proxy for presynaptic strength variability, with lower PPR disparity indicating lower PPR variability and vice versa (Figure S5A,B). Consistent with our analysis thus far, bath application of MK801 and AP5 reduced PPR disparity (Figure S5D,E; MK801 p = .001, AP5 p=.004), while Ro25-6981 and MPEP did not (Figure S5G,H; Ro25-6981 p = .670, MPEP p = .263).

Collectively, these findings support the idea NMDARs play a role in bi-directionally regulating presynaptic strengths to broaden the variability of PPR. Moreover, such an involvement of NMDARs in presynaptic regulation is distinct from the presynaptic regulation by mGluR5 that potentiates synaptic strength across a synapse population.

### Astrocyte NMDARs mediate the effects of NMDAR antagonists on PPR diversity

To test the extent to which bath applied NMDAR antagonists affected synaptic transmission by targeting astrocyte NMDARs, we knocked down in astrocytes, the *GRIN1* gene that encodes GluN1, a requisite subunit for the cell surface expression of NMDARs (Fukaya et al., 2003; Abe et al., 2004). AAV carrying either mCherry-tagged nlsCre (Cre) or a control nls-mCherry lacking Cre (ΔCre) under the GFAP104 promoter was injected bilaterally into the dorsal hippocampus of adult *GRIN1* floxed mice (Figure 2A)(Letellier et al., 2016). We used low titer AAVs (∼0.4-4E10 genome copies/injection) to avoid potential reactive astrocytosis, and although the extent was modest, we obtained a highly specific mCherry expression in astrocytes throughout the area CA1 with little expression in neurons (Figure 2B: Cre, 41.1 ± 2.4% of GFAP+ve cells [n = 3529], 0.3 ± 0.1% of NeuN+ve cells [n = 1150] from 5 mice; ΔCre, 40.7 ± 1.9% of GFAP+ve cells [n = 3309], 0.5 ± 0.1% of NeuN+ve cells [n = 1217] from 5 mice). The efficacy of GluN1 knock-down in astrocytes was assessed electrophysiologically by patch-clamping astrocytes and monitoring slow depolarizing responses elicited by puff applying NMDA and glycine (1mM each) (Figure S6)(Letellier et al., 2016). Astrocytes that expressed Cre as identified by the mCherry fluorescence in *stratum radiatum* (SR) where Schafer collateral synapses were present, showed depolarizing responses to NMDA-glycine puff that were significantly decreased compared to ΔCre controls (Figure S6A; p = 0.006). Given that mCherry-positive astrocytes were present throughout the CA1, we also monitored NMDA-glycine puff responses in astrocytes in *stratum oriens* (SO) and *stratum lacunosum moleculare* (SLM). In contrast to mCherry-positive astrocytes in SR, mCherry-labelled astrocytes in SO or SLM showed depolarizing responses to NMDA-glycine puff that were not different between Cre and control ΔCre slices (Figure S6B-C; SO p = .247; SLM p = .268), despite the fact that viral transduction efficiencies were comparable amongst astrocytes across the three layers (SR: Cre 39.8 ± 2.4%, ΔCre 40.2 ± 2.0%; SO: Cre 44.8 ± 3.0%, ΔCre 39.3 ± 5.9%; SLM: Cre 39.1 ± 4.9%, ΔCre 40.0 ± 4.0%). The input resistance of patched astrocytes was not altered by Cre expression in any layer (Figure S6D). Notably NMDA-glycine responses in SLM were substantially smaller than those in SR or SO (p = 0.001, one way ANOVA, Bonferroni post-hoc tests). Nevertheless, AP5 bath application blocked NMDA-glycine puff-induced depolarizing responses in astrocytes in all three layers. This observation suggests that NMDARs are broadly expressed across hippocampal CA1 astrocytes. Moreover, the differential sensitivity of the slow depolarizing responses to astrocyte GluN1 knock-down between layers supports the possibility that the cumulative actions of signaling downstream to NMDAR activation including the contribution of voltage-gated conductances (Serrano et al., 2008; Letellier et al., 2016; Shih et al., 2013) could be heterogeneous across layers. We will return to this layer-specific differences later.

**Figure 2.**
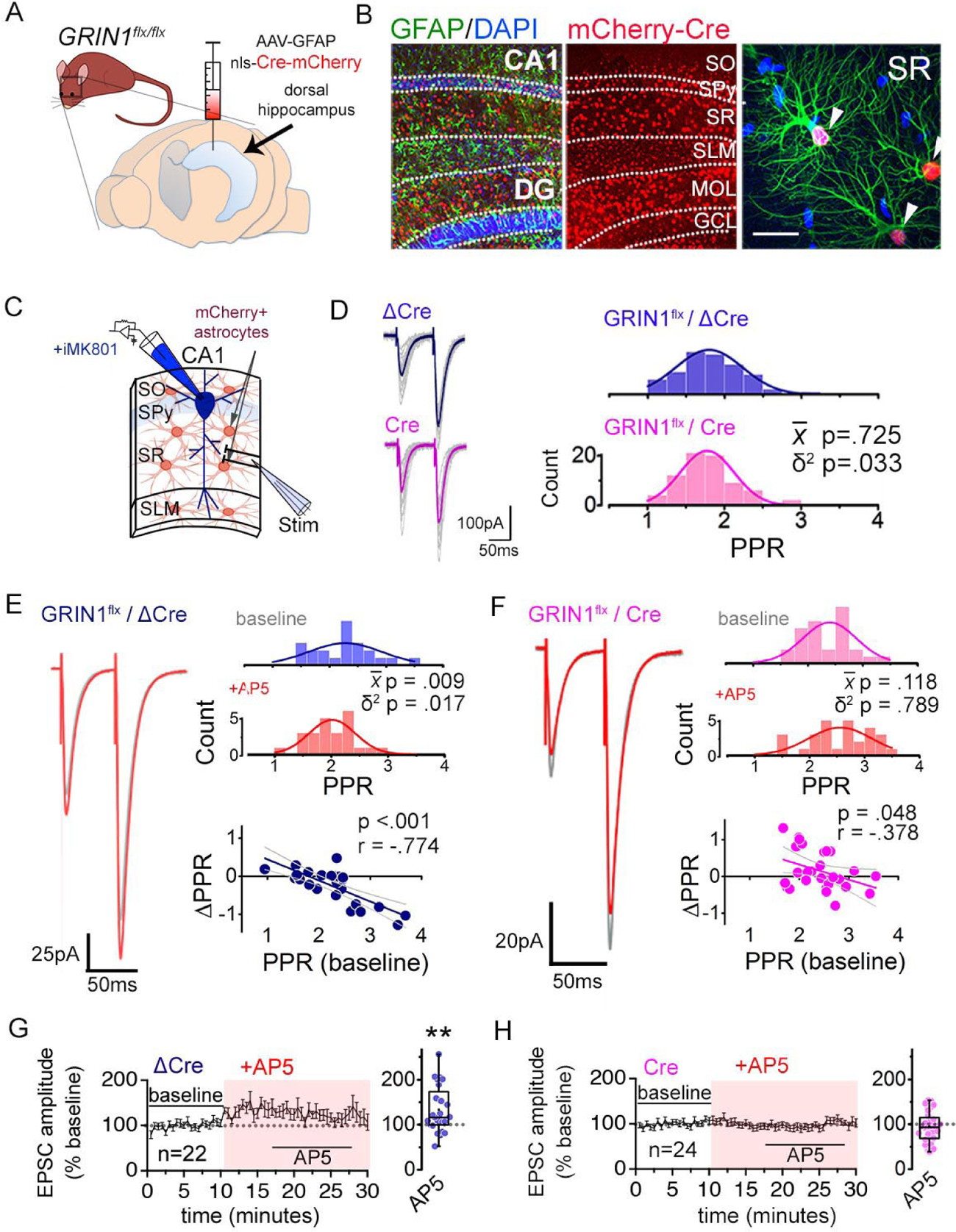
Astrocyte NMDARs mediate the antagonist-induced decrease in synaptic strength diversity. (**A**) Astrocyte GluN1 was conditionally knocked down by injecting AAV-GFAP104-nls-mCherry-Cre (Cre), or as a control, AAV-GFAP104-nls-mCherry (ΔCre), to the dorsal hippocampus of adult GRIN1*^flx/flx^* mice. (**B**) Representative image of hippocampal area CA1 immunolabelled for GFAP (green) and mCherry (red) and counterstained with DAPI (blue), 21 days after virus injection. SO, *stratum oriens*; SPy, *stratum pyramidale*; SR, *stratum radiatum*; SLM, *stratum lacunosum moleculare*; MOL, molecular layer; GCL, granule cell layer. Right: magnified view in SR showing restricted expression of mCherry to GFAP-positive astrocytes (arrowheads). Scale bar, 20 µm. (**C**) Recording scheme: CA1 pyramidal neurons were patched in acute slices prepared 21 days after ΔCre or Cre AAV infection. EPSCs were elicited by placing the stimulating electrode in the SR. Patch pipette contained MK801 for experiments in *E-H*. (**D**) Left: Averaged EPSC traces from representative recordings in slices from control ΔCre (top) or Cre (bottom) virus-infected mice. Individual traces are shown in grey. Right: Histograms of PPRs recorded from ΔCre (top) and Cre slices (bottom) were fit with single Gaussians. The p values were obtained for the population mean (x̅) (two-tailed t-test) and variance (δ^2^) (one-tailed f-test for equal variances) comparing PPR distributions from ΔCre (n=72 inputs, 43 cells, 14 mice) and Cre (n=74 inputs, 46 cells, 12 mice) slices. (**E,F**) Left: average EPSC traces in baseline (grey) and after drug treatment (color) from a representative experiment. Right: PPR histograms (top) and ΔPPR vs. baseline PPR plots (bottom) in baseline and after bath applying AP5 (50 µM) to ΔCre (*E*) and Cre (*F*) slices. The p values for the population mean (x̅) and variance (δ^2^) are as in *D*, comparing PPRs before and after the drug treatment. Linear fits and Pearson’s correlation coefficients (r) and p values are shown in scatter plots. (**G,H**) Plots of normalized EPSC amplitudes before and during the application of AP5 to ΔCre (*G*) and Cre-infected (*H*) slices (shaded area); n is the number of inputs examined. Right: summary bar graph. **p = 0.005, Mann-Whitney U-test. ΔCre +AP5 = 22 inputs, 12 cells, 5 mice; Cre +AP5 = 24 inputs, 14 cells, 5 mice. (Figure 2 legend continued from previous page)

Having detected GluN1-dependent functional NMDAR activity in SR CA1 astrocytes, we next asked whether the GluN1 knock-down also affected synaptic transmission of Schaffer collateral synapses similarly to the bath applied NMDAR antagonists. Baseline PPR showed a significantly reduced PPR variance in Cre-infected slices compared to control slices without a change in the mean PPR (Figure 2C,D; δ^2^ p = .033; x̅ p = .725). Analysis of the relative PPR from two independent Schaffer collateral inputs also revealed lower PPR disparity in Cre injected slices compared to control slices (Figure S7A; p = .009). These observations are consistent with a reduced PPR variance upon compromising astrocyte NMDAR signaling as observed with bath applied NMDAR inhibitors. The amplitude and frequency of spontaneous EPSCs and the waveform of evoked EPSCs were unaltered by astrocyte-specific GluN1 knock-down (Figure S7B,C).

We next determined whether astrocyte GluN1 knock-down could occlude the effects of bath applied AP5 on PPR variance, by recording from postsynaptic CA1 neurons infused with MK801 in ΔCre and Cre-infected slices. Similar to naïve slices, in ΔCre control slices, AP5 decreased the population variance of PPRs by strengthening some synapses and weakening others (Figure 2E; ΔCre +AP5 p = .017). In addition, AP5 caused also a variable increase in EPSC amplitudes (Figure 2G; AP5 p = .005) that occurred without a concomitant change in the amplitude or the frequency of spontaneous EPSCs (Figure S7D,E). In Cre slices, however, the decrease in PPR variance by AP5 was strongly attenuated, despite the modest extent of virus infection in Cre slices (Figure 2F; Cre +AP5 p = .789). The analysis of PPR disparity across two inputs also showed substantially decreased inhibitory effect of AP5 in Cre slices relative to controls (Figure S7H,I; ΔCre +AP5 p = .013, Cre +AP5 p =.315). Curiously, the small increase in EPSC amplitudes observed upon applying AP5 was also attenuated in Cre slices (Figure 2H). This suggests that although the effect of NMDAR antagonists on PPR diversity and EPSC amplitudes are likely to target distinct compartments – presynaptic and postsynaptic – both mechanisms may involve astrocyte NMDARs. Together, these observations indicate that astrocyte NMDARs are the major mediators of the synaptic effects of acutely blocking NMDARs in our experimental conditions, and further support the view that astrocyte NMDARs help maintain the broad variability of presynaptic strengths across a synapse population.

### Modelling of release probability variability reveals its impact on synaptic plasticity

In order to gain an insight into the physiological role for the broad release probability distribution, we sought to assess the impact of reducing the presynaptic strength variability on a synaptic learning rule by constructing a mathematical model. We first modelled release probability variations based on log-normal distributions (Buzsáki and Mizuseki, 2014; Branco and Staras, 2009; Murthy et al., 1997) for a range of peak release probability distributions (*U*_max_) that represented the control condition and when the variance was reduced to mimic conditions of astrocyte NMDAR blockade, termed “control” and “NMDAR blocked”, respectively (Figure 3A; see Methods). The optimal parameters for the log-normal distributions were obtained from the best fit to a modified synaptic facilitation model based on Tsodyks et al. (1998) using the experimental data in which the peak EPSC amplitudes to repetitive stimulation of the Schaffer collaterals at 20 Hz were monitored under identical conditions to the PPR experiments (Figure S8A; see Methods). Next, we used the BCM learning rule (Bienenstock et al., 1982) to investigate potential implications of altering the variance of release probability on long-term synaptic plasticity. In the BCM learning rule, the synaptic weight undergoes changes according to the excitatory postsynaptic current and the presynaptic neurotransmitter release, in which the magnitude and the type of long-term plasticity can be controlled by a set of parameters (see Methods). In simulations, we explored the effect caused by decreasing the variance of the release probability on both long-term potentiation (LTP) and long-term depression (LTD). Figure 3B illustrates an example experiment for *U*_max_ = 0.1, where traces of the synaptic weight function w(t) for LTP and LTD are two conditions that differ only with respect to the release probability variance of the presynaptic neuron. Application of 100 presynaptic spikes with 50-ms inter-spike intervals triggered a rapid change in synaptic weight during the stimulation that was stably maintained after the stimulation for both LTP and LTD (Figure 3B). Although the efficacy of both LTP and LTD are reduced in the condition of less variable release probability mimicking the NMDAR-block situation, the effect is larger for LTP.

**Figure 3.**
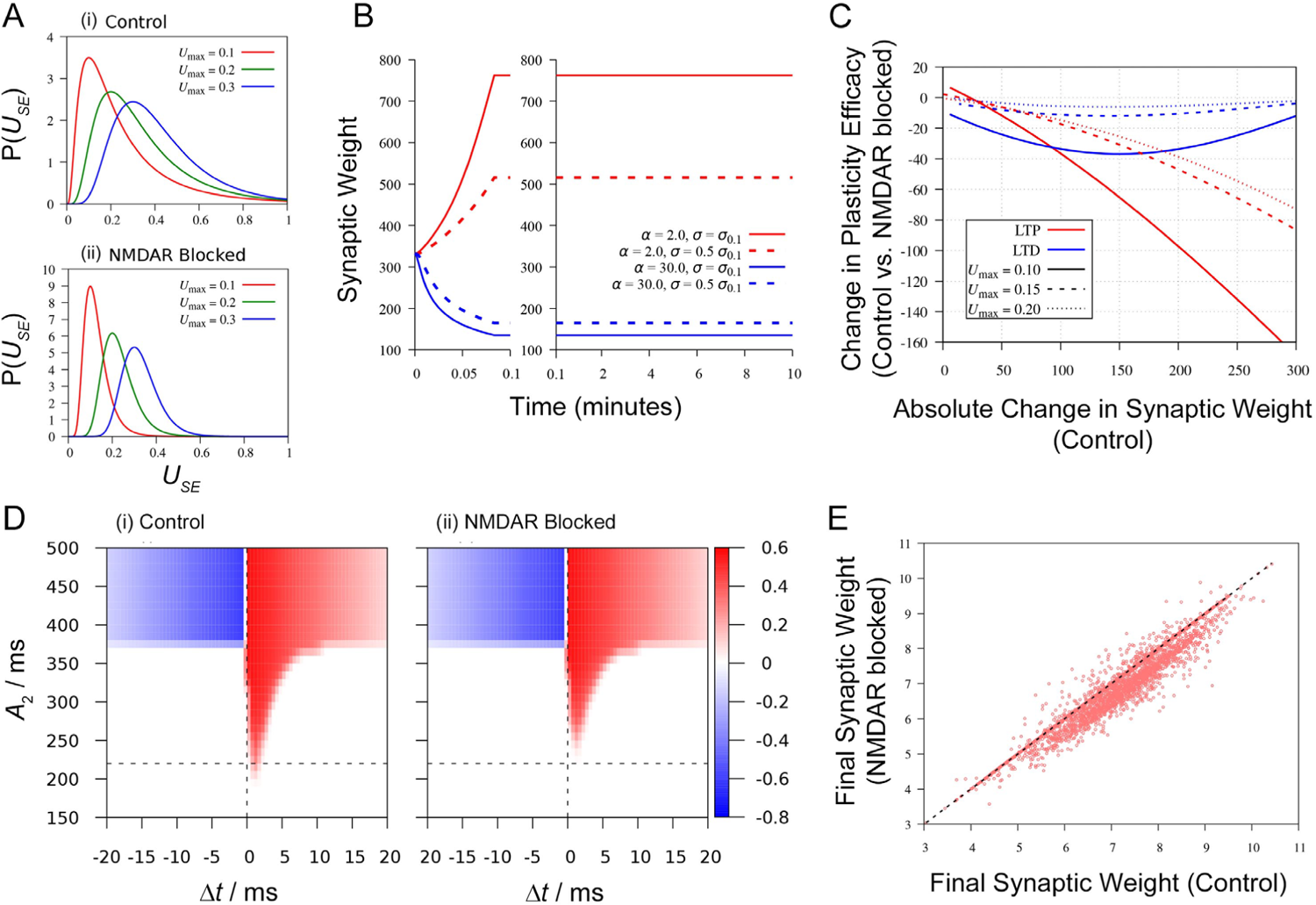
Synaptic strength diversity impacts long-term synaptic plasticity. (**A**) Variations of release probability distribution *U*_SE_, with different peak locations (*U*_max_ = 0.1, red; *U*_max_ = 0.2, green; *U*_max_ = 0.3, blue) modelled using a log-normal distribution reported by Murthy et al., (1997). The plot shows release probability distributions for the control condition (i) and the condition of reduced release probability variance mimicking the NMDAR-block condition (ii). (**B**) Example results of the BCM-like learning simulation. Traces show synaptic weight function w(t) for LTP (red: α = 2.0) and LTD (blue: α = 30.0) simulations where long-term synaptic plasticity is induced by applying 100 presynaptic spikes with 50 ms inter-spike intervals; the decay of synaptic plasticity has not been considered in the model. Two limiting cases of the release probability distributions of the presynaptic neuron (control, solid line; NMDAR block, dotted line; *U*_max_ = 0.1 for both cases) as illustrated in (*Ai*) and (*Aii*) are shown. (**C**) Plot of the difference of the synaptic weight change at *w*(t = 2 min) between the release probability distributions representing control and NMDAR block for LTP (red) and LTD (blue) as a function of the change in synaptic weight *w*(t = 2 min) for the control case over a range of potentiation or depression. The horizontal axis is defined by *w(t = 2 min, σ = σ_max_) − w(t = 0)*. The difference of synaptic weight in the y-axis is defined as w(t = 2 min, σ = σ_max_/2) − w(t = 2 min, σ = σ_max_) for LTP and as w(t = 2 min)-w(t = 2 min, σ = σ_max_/2), for LTD. The relationship is shown for three different peak distributions, *U*_max_, for both LTP and LTD (*U*_max_ = 0.1, solid line; *U*_max_ = 0.2, dashed line; *U*_max_ = 0.3, dotted line). (**D**) Simulation result using two leaky integrate-and-fire neurons. Changes in synaptic weight *w*(t) after giving 10 spike pairs in which the current injection to the presynaptic neuron is 500 pA and the postsynaptic neuron is *A*_2_, in control (i) and in NMDAR blocked condition (ii). The current injection sequences to the presynaptic and the postsynaptic neurons are separated by a time difference Δ*t*. Color intensity shows potentiation (red) and depression (blue) of synaptic weight changes. Vertical dashed line: Δ*t* = 0. Horizontal dashed line: the least *A*_2_ to evoke LTP under the NMDAR-block condition. (**E**) A comparison of the simulation result using Poisson spike trains. Each data point represents the synaptic weights after applying the same pair of Poisson sequences for the release probability distribution representing the control condition (x-axis) and the NMDAR block condition (y-axis). The NMDAR block condition consistently shows smaller synaptic weight compared to the control condition. (Figure 3 legend continued from previous page)

We next sought to test whether the qualitative result of the simulation is preserved over a range of LTP and LTD (i.e. for a series of parameter *α*; see Methods). To this end, the difference of synaptic weight change induced by stimulation between conditions of broad (control) vs. narrow (NMDAR block) release probability variability was considered as a function of a range of the absolute change in synaptic weight w(t) under control release probability conditions for both LTP and LTD (Figure 3C, solid lines). In the numerical exploration, we found that LTD was always smaller if the pre-synaptic neuron had less variable release probability. In contrast, the sensitivity of LTP to release probability variance depended on the magnitude of potentiation, in which the adverse effect of the narrowing of release probability distribution in compromising synaptic plasticity was increasingly pronounced with a larger degree of potentiation. The robustness of the observed sensitivity of LTP and LTD to the differences in release probability variance was further tested by changing the peak release probability *U*_max_ under the same conditions (*U*_max_ = 0.10, 0.15, 0.20). The overall pattern of the decrease in the efficacy of LTP and LTD persisted upon narrowing the release probability distribution. However, the reduction of LTD became less prominent when *U*_max_ was increased (Figure 3C).

In order to determine whether the influence of release probability variance on the efficacy of long-term plasticity could be observed in a different model, spike-timing dependent plasticity (STDP) was simulated using synaptically connected two leaky integrate-and-fire (LIF) neurons (see Methods). The presynaptic neuron received a spike train of 500 mA magnitude, *A*1, while the postsynaptic neuron received a spike train of variable magnitude *A*_2_. Pairing 10 spikes at 20 Hz with a time difference between spike trains of Δ*t* resulted in LTP or LTD, with positive Δ*t* values producing LTP and negative Δ*t* values resulting in LTD (Figure 3D(i)). Notably, conditions that mimic decreased release probability variance seen in NMDAR-blocked experiments showed no observable changes in LTD whereas LTP was compromised compared to control conditions (Figure 3D(ii)), the latter effect on LTP being similar to the observations in the BCM-like model.

Finally, the impact of narrowing the release probability variance was explored under a more general STDP scenario by examining synaptic weight changes elicited in a pair of neurons receiving Poisson spike trains. Specifically, the synaptic weights obtained in response to the same Poisson sequences in the control condition or the NMDAR-blocked condition for a number of independent simulations at *U*_max_ of 0.10 showed that the efficacy of potentiation was consistently smaller in the NMDAR-blocked condition (Figure 3E). Such an attenuating effect of the reduced release probability variance on potentiation was observed also when the peak of release probability distributions was shifted to larger values (*U*_max_ = 0.15 and 0.20; Figure S8C).

Altogether, the numerical simulation experiments demonstrate that a reduced variability in presynaptic release probability can reshape the long-term synaptic plasticity dynamics. In the BCM-like model, both LTP and LTD were reduced, and in the LIF-neuron-pair model, LTP was reduced whereas LTD appeared largely unchanged. Collectively, LTP is likely to be compromised across a synapse population when the distribution of release probability is narrowed, while the effect on LTD could depend on the fine detail of the spike sequence.

### Hippocampal astrocytes express GluN2C NMDARs

Our results thus far point to astrocyte NMDARs, and in particular, the involvement of the obligatory subunit GluN1 in regulating the variability of presynaptic strengths. In order to obtain direct molecular evidence for the expression of NMDAR in hippocampal astrocytes and to clarify the relevant NMDAR subtype, we performed single cell RT-PCR to compare the expression of GluN1, GluN2A, GluN2B and GluN2C mRNAs in CA1 astrocytes across SO, SR and SLM layers and CA1 pyramidal neurons (Figure 4A). Acute hippocampal slices were prepared from adult mice as used for electrophysiology experiments, and after labelling astrocytes with sulforhodamine101 (Nimmerjahn et al., 2004), RNA from single cells was extracted by patch-clamping. All putative astrocytes had low input resistances (SR 11.20 ± 0.36 MΩ; SO 13.24 ± 0.43 MΩ; SLM 10.60 ± 0.33 MΩ; n = 39 cells from 9 mice sampled for each layer), lacked action potentials, and displayed linear current-voltage relationships (Figure 4A,B). CA1 pyramidal neurons expressed high levels of GluN1, GluN2A and GluN2B subunit mRNAs but expressed substantially low levels of GluN2C mRNA. In contrast, astrocytes in all three layers showed robust expression of GluN2C mRNA. Moreover, astrocytes showed minimal expression of GluN2A and GluN2B mRNAs, and the GluN1 mRNA expression in astrocytes was also substantially low compared to neurons (Figure 4C). GluN2D mRNA in CA1 pyramidal neurons and astrocytes was also tested, and it was undetectable in RNA extracted from patch clamping both cell types while the probe itself could detect GluN2D mRNA in brain tissue extracts (data not shown). Inability to detect GluN2D mRNA in astrocytes is consistent with previous reports of single cell RNA-seq analysis (Khakh Lab database; also see Alsaad et al., 2019). We also examined the levels of expression of NMDAR subunits in CA3 pyramidal neurons, and the pattern of subunit expression mirrored the pattern observed for CA1 pyramidal neurons (data not shown).

**Figure 4.**
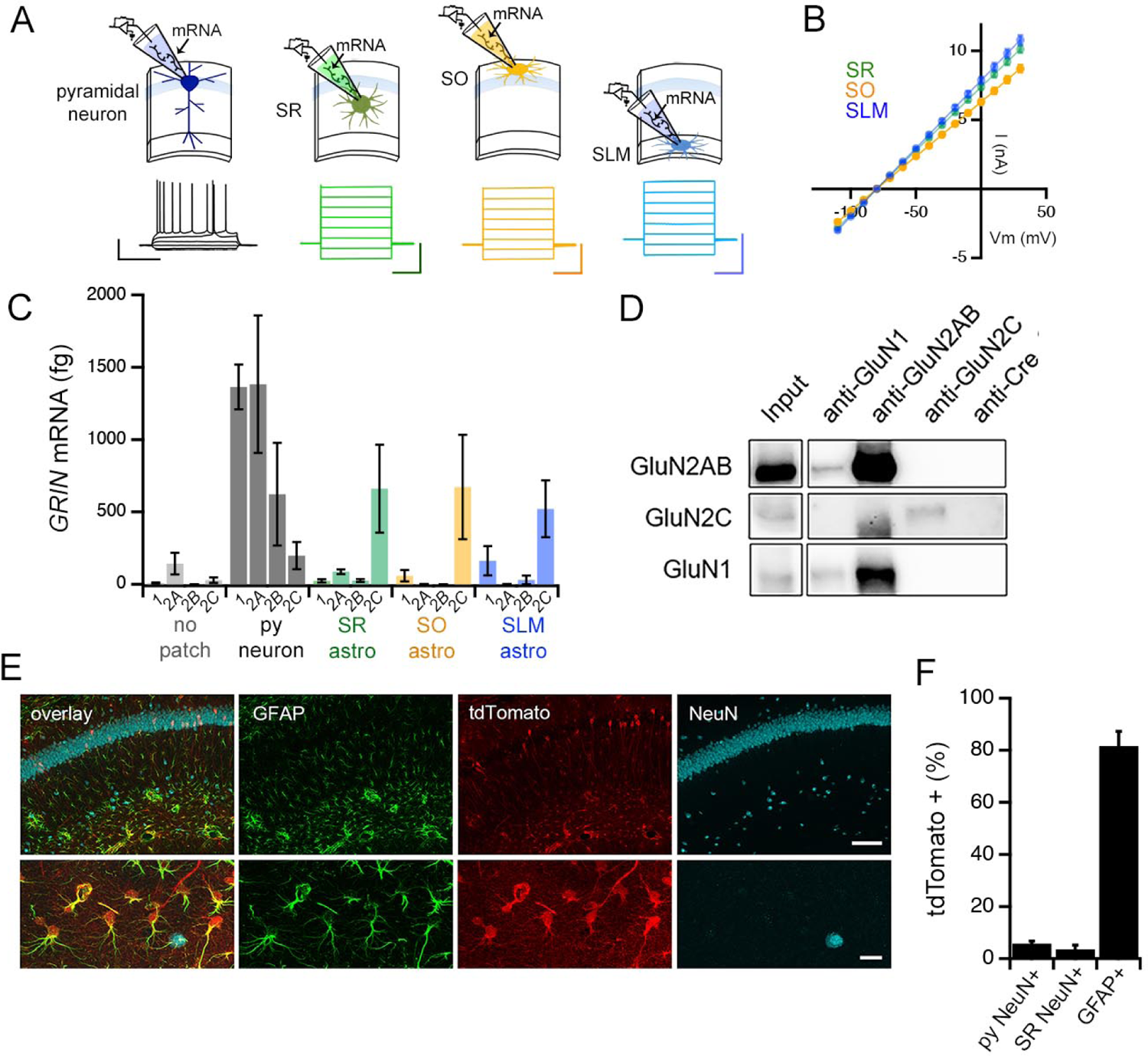
Adult mouse hippocampal astrocytes express GluN2C-NMDA receptors. (**A**) Top: patch RT-PCR strategy. Bottom: representative voltage responses to current steps in CA1 pyramidal neuron (far left, black), and current responses to voltage steps in astrocytes in *stratum radiatum* (SR, green), *stratum oriens* (SO, yellow), and *stratum lacunosum moleculare* (SLM, blue). Scale bars: neuron, 40 mV, 500 ms; astrocytes, 6 nA, 200 ms. (**B**) Summary of I-V curves of astrocytes obtained before collecting RNA. N = 39 cells, 36 slices, 9 mice for each layer. (**C**) Summary of RT-PCR analysis for mRNA encoding indicated NMDAR subunits: *GRIN1* (1), *GRIN2A* (2A), *GRIN2B* (2B), and *GRIN2C* (2C). Control samples (no patch) were obtained by inserting the electrode into the slice without patching cells. *GRIN2D* levels were undetectable in all cells examined (data not shown). (**D**) Western blots of co-immunoprecipitations performed using adult mouse hippocampal extracts with the indicated antibodies where anti-Cre antibody is used as a negative control. The blots are probed for GluN2A/2B (top row), GluN2C (middle row) or GluN1 (bottom row). (**E**) Representative brain sections of *GRIN2C*-Cre mice crossed to a tdTomato reporter line (Ai9), showing hippocampal area CA1 immunolabelled for GFAP, tdTomato and NeuN; scale bars, 160 μm (top), 25 μm (bottom). (**F**) Quantification of % cells that are positive for tdTomato amongst NeuN-labelled cells in *stratum pyramidale* (left) and SR (middle) and GFAP-labelled cells (right).

The expression of GluN2C subunit protein in the hippocampus was confirmed by immunoprecipitation of mouse hippocampal extracts followed by Western blotting, using an antibody against the Cre recombinase that is not expressed in wild type mice, as a negative control (Figure 4D). For characterizing the cellular expression pattern of GluN2C in the hippocampus, we were unable to identify an antibody against GluN2C that was suitable for immunofluorescence labelling experiments. Therefore, in order to visualize the cells that expressed GluN2C, we used a GluN2C mutant mouse line carrying an insertion of a codon-improved Cre recombinase immediately downstream of the translation initiation site in the *GRIN2C* gene (Miyazaki et al., 2012). GluN2C-Cre mice were crossed with a Cre reporter line (Ai9: Madisen et al., 2010) that expressed tdTomato upon Cre-mediated recombination. The hippocampus of the offspring showed robust tdTomato fluorescence in astrocytes identified by GFAP labelling (81.6 ± 5.7% of GFAP+ve cells; Figure 4E,F), which was consistent with the high expression level of GluN2C mRNA in astrocytes (Figure 4C). Moreover, some interneurons in SR identified by the NeuN labeling also expressed tdTomato (3.6 ± 1.6% of NeuN+ve cells in SR, Figure 4E,F), which was in agreement with previous reports (Ravikrishnan et al., 2018; Gupta et al., 2016). Surprisingly, the pyramidal cell layer also showed sporadic tdTomato-labelled neuronal cell bodies, although such cells represented a limited proportion of pyramidal neurons (5.8 ± 1.0% of NeuN+ve cells in *stratum pyramidale*, Figure 4E,F). The RT-PCR, biochemical and immunohistochemistry experiments collectively provide evidence in support of GluN2C-containing NMDAR as a major NMDAR type expressed in hippocampal CA1 astrocytes.

### Inhibition of GluN2C NMDARs reduces PPR variability in a manner sensitive to astrocyte GluN1 expression

Given the prominent expression of GluN2C in astrocytes, we wondered if the regulation of PPR could be attributed to GluN2C NMDARs. To test such a possibility, we examined whether pharmacological inhibition of GluN2C NMDARs could mimic the effects of bath applied AP5 and MK801 on PPR variance using QNZ46 (25 µM), an NMDAR antagonist specific for GluN2C/D (Hansen and Traynelis, 2011). Similarly to MK801 and APV, QNZ46 reduced the population variance of PPRs without altering the mean PPR (Figure 5A; δ^2^ p=.008; x̅ p=.312). Again we observed the negative correlation between the change in PPR and the baseline PPR (Figure 5A, bottom), with a linear fit to the data intercepting the x-axis near the baseline mean PPR (X intercept = 2.07). Additionally, in experiments monitoring two independent Schaffer collateral inputs, bath application of QNZ46 decreased the PPR disparity as found for AP5 and MK801 (Figure S5F; p = .005). These observations suggest that GluN2C/D-containing NMDARs contribute to maintaining the broad PPR diversity. Given the lack of detectable expression of GluN2D in hippocampal CA1 cells, it is likely that NMDARs containing GluN2C mediate the observed effects of QNZ46 in modulating presynaptic strength to decrease the overall range of PPR variability.

**Figure 5.**
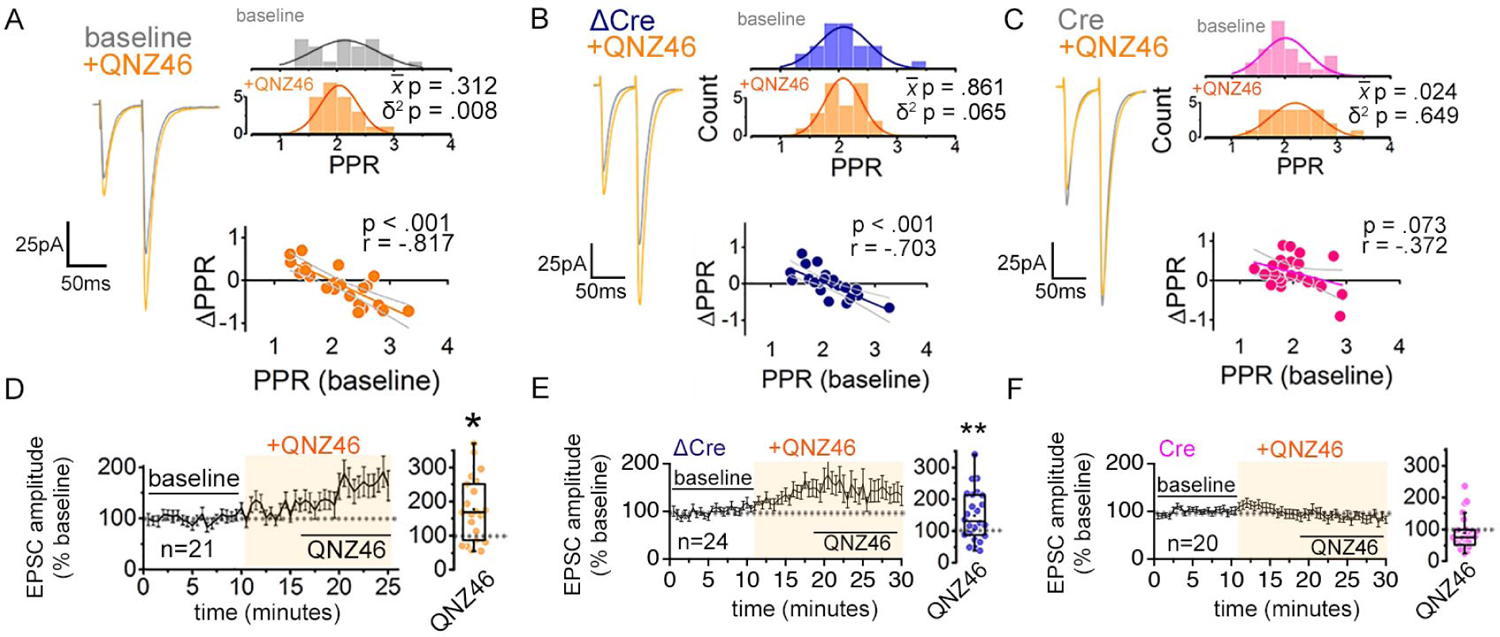
Inhibiting GluN2C NMDARs reduces PPR variability in a manner dependent on astrocyte expression of GluN1. (**A-C**) Left: representative EPSC traces (average of 20 sweeps) to pairs of pulses (Δt = 50 ms) applied to Schaffer collateral axons in baseline and after wash-on of QNZ46 (25 µM). Right: PPR histograms (top) and ΔPPR vs. baseline PPR plots (bottom) before and after bath applying QNZ46 to control uninfected (*A*), ΔCre (*B*) and Cre (*C*) slices. The p values were obtained for the population mean (x̅) (paired sample t-test) and variance (δ^2^) (one-tailed f-test for equal variances) comparing PPRs before and after the drug treatment. Linear fits and Pearson’s correlation coefficients (r) and p values are shown in scatter plots. (**D-F**) Plots of EPSC amplitudes (normalized to the baseline average) before and during the application of QNZ46 to control uninfected (*D*), ΔCre (*E*) and Cre-infected (*F*) slices (shaded area) where n is the number of inputs examined for each experiment. Baseline and experimental periods are indicated (black bars). Box plots show the summary of normalized EPSC amplitudes during the QNZ46 application. *p<0.05, **p = 0.005, Mann-Whitney U-test was used to compare the control to the drug conditions. No virus infection +QNZ46 = 21 inputs, 11 cells, 7 mice; ΔCre +QNZ46 = 24 inputs, 12 cells, 5 mice; Cre +QNZ46 = 20 inputs, 12 cells, 6 mice.

When EPSC amplitudes were examined, QNZ46 bath application caused a significant increase, reminiscent of the changes observed upon AP5 and MK801 application (Figure 5D; p = .014). Moreover, similarly to AP5, QNZ46 significantly increased CV^-2^ of EPSC amplitudes (Figure S3B; p = .009), which was suggestive of presynaptic alterations.

Next we asked whether QNZ46 acted on astrocyte NMDARs to control PPR variability. To address this point, we again tested for occlusion using astrocyte GluN1 knock-down slices. QNZ46 was bath applied to Cre and ΔCre-infected slices while recording EPSCs from CA1 neurons infused with MK801. In ΔCre control slices QNZ46 decreased the PPR variance (Figure 5B; p = .065) and caused a variable increase in EPSC amplitudes (Figure 5E; p = .005) without a concomitant change in the amplitude or the frequency of spontaneous EPSCs (Figure S7F). Crucially, as observed for AP5, QNZ46 was no longer effective in producing a significant change in the PPR variance in Cre slices (Figure 5C; p = .649). When PPR disparity across two inputs was monitored, QNZ46 decreased the PPR disparity in a manner that was sensitive to astrocyte GluN1 knock-down, which also supported the involvement for GluN2C-NMDARs in regulating presynaptic strength diversity (Figure S7J,K; ΔCre +QNZ46 p = .031; Cre +QNZ46 p = .126). Moreover, similarly to AP5, QNZ46-dependent increase of EPSC amplitudes was attenuated also in Cre slices (Figure 5F).

Together, these observations indicate that GluN2C NMDARs expressed in astrocytes function to maintain the broad variability of presynaptic strengths across a synapse population. While GluN2C NMDARs also play a role in regulating EPSC amplitudes, this is likely to involve a mechanism that is distinct from the presynaptic regulation but is engaged in parallel to target postsynaptic processes.

### Layer-specific regulation of synaptic strength by astrocyte NMDARs

Astrocytes are a highly heterogeneous cell type (Zhang and Barres, 2010; Khakh and Sofroniew, 2015) that influence synaptic transmission in a synapse- and circuit-specific manner (Martin-Fernandez et al., 2017; Schwarz et al., 2017; Martin et al., 2015; Lanjakornsiripan et al., 2018). Our RT-PCR analysis of NMDAR subtype expression showed relatively high expression of GluN2C mRNA across SR, SO and SLM layers in CA1 astrocytes compared to CA1 pyramidal neurons. Nevertheless, despite the relatively even expression of GluN2C across layers, our NMDA-glycine puff application experiments CA1 astrocytes revealed layer-specific differences in the GluN1-dependent component of the slow depolarizing responses triggered by the NMDA-glycine puff, with SR astrocytes showing the highest sensitivity to the astrocyte GluN1 knock-down (Figure S6). This raised the possibility that astrocyte NMDAR-dependent modulation of synaptic inputs to CA1 pyramidal neurons whose dendrites span across SR, SO and SLM, might also differ across layers.

To test such possibility of layer-specific regulation, we compared astrocyte NMDAR-dependence of synaptic transmission in CA1 neurons across three layers in slices from astrocyte GluN1 knock-down and control mice. In contrast to the specific decrease in the PPR variance observed in Cre slices relative to ΔCre control slices for the SR input (Figure 2D), neither the PPR variance nor its mean differed between Cre and ΔCre control slices for the SO and the SLM inputs (Figure 6A-D; SO: δ^2^ p = .303, x̅ p = .402; SLM: δ^2^ p = .624, x̅ p = .161). We also examined the layer-specificity of PPR diversity regulation using a pharmacological approach in slices from wild type mice. The sensitivity of PPRs to bath applied NMDAR antagonists in CA1 pyramidal neurons intracellularly perfused with MK801, was monitored in response to pairwise stimulation of SO and SLM inputs as had been done for the SR input. Neither MK801 nor AP5 had any appreciable effect on PPR variance in SO (Figure 6E,G: MK801, p = .107; AP5 p = .472) and SLM (Figure 6I,K: MK801, p = .920; AP5, p = .741). MK801 modestly increased EPSC amplitudes at some inputs in SO (Figure 6F: p = .783) but it had no effect in SLM (Figure 6J: p = .915). Moreover, AP5 did not potentiate EPSC amplitudes as observed for the SR input, but instead it significantly depressed EPSC amplitudes in SO (Figure 6H: p = .028) while it caused no appreciable change in SLM (Figure 6L: p = .317). Collectively, these results suggest that regulation of PPR diversity by astrocyte NMDARs in CA1 pyramidal neurons is specific to the SR inputs.

**Figure 6.**
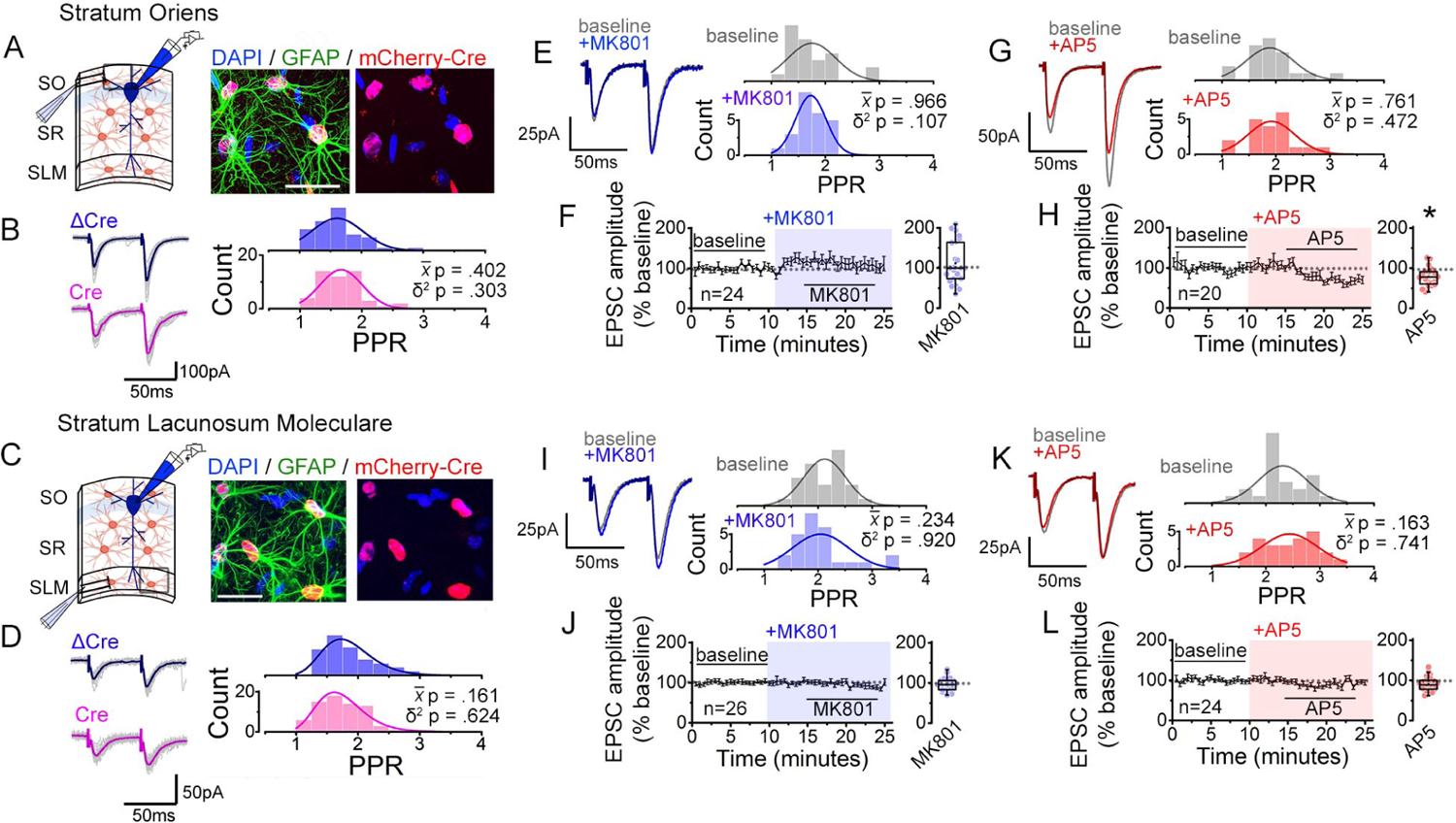
Interfering with astrocyte NMDARs has little effect on synaptic strength diversity in the *stratum oriens* (SO) and the *stratum lacunosum moleculare* (SLM). (**A**) Left: recording scheme. Right: confocal image from SO in CA1 showing nls-mCherry-Cre-positive astrocytes labelled with GFAP and DAPI. Scale bar, 20 µm. (**B**) Average (color) and individual (grey) EPSC sweeps from representative recordings by stimulating SO of ΔCre and Cre infected slices. PPR histograms from ΔCre (top) and Cre slices (bottom) fit with single Gaussians. [p values: comparisons of population mean (x̅) (Mann-Whitney test) and variance (δ^2^) (one-tailed f-test for equal variances) between ΔCre and Cre.] ΔCre, n = 50 inputs, 25 cells, 7 mice; Cre, n = 50 inputs, 25 cells, 7 mice. (**C,D**) Same analysis as in *A,B* for SLM. ΔCre, n = 67 inputs, 47 cells, 12 mice; Cre, n = 71 inputs, 49 cells, 12 mice. (**E,G**) Representative EPSC traces and PPR histograms of from SO stimulation in baseline and after applying MK801 (*E*) or AP5 (*G*). (**F,H**) Normalized EPSC amplitudes before and after applying MK801 (*F*) or AP5 (*H*) (shaded area) in SO. Right, summary bar graph. MK801, n = 24 inputs, 12 cells, 6 mice; AP5, n = 20 inputs, 10 cells, 7 mice. * p<0.05 Mann-Whitney U-test. (**I,K**) Same analysis as in *E,G* for PPRs elicited in SLM in control and MK801 (*I*) or AP5 (*K*). (**J,L**) Same analysis as in *F,H* with MK801 (*J*) or AP5 application (*L*). MK801: n = 26 inputs, 13 cells, 8 mice; AP5, n = 24 inputs, 12 cells, 6 mice.

As a further test of the layer specificity of astrocyte NMDAR-dependent regulation of PPR variability, we assessed PPR disparity by performing two independent pathway analysis in each layer in response to bath applied NMDAR antagonists. Again, in contrast to SR input (Figure S5D, E), neither MK801 nor AP5 depressed the PPR disparity of the SO and the SLM inputs (Figure S9). These observations are consistent with the role for astrocyte NMDARs, and particularly GluN2C NMDARs, for maintaining the diversity of PPR specifically of SR inputs to CA1 pyramidal neurons.

## Discussion

Broad heterogeneity in the efficacy of synaptic transmission is a fundamental feature of small glutamatergic synapses in the mammalian brain (Dobrunz and Stevens, 1997; Atwood and Karunanithi, 2002; Branco and Staras, 2009). A variety of presynaptic parameters contribute to the observed variability, such as the abundance, subtype, and location of presynaptic calcium channels (Thalhammer et al., 2017; Brenowitz and Regehr, 2007; Eltes et al., 2017), their positioning with respect to the synaptic vesicles, the active zone size and the number of docked vesicles, the state of vesicle fusion machinery (Park and Tsien, 2012; Marra et al., 2012; Fulterer et al., 2018; Glebov et al., 2017; Holderith et al., 2012), as well as the coupling to neuromodulatory signals (Burke et al., 2018), and the synapse’s recent history of synaptic plasticity. These determinants of presynaptic release efficacy are influenced by target-specific signals (Éltes et al., 2017; Branco et al., 2008; Reyes et al., 1998; Markram et al., 1998) and are further subject to activity-dependent modulation (Thalhammer et al., 2017; Goda and Stevens, 1998; Burke et al., 2018). Although the basis for presynaptic release probability regulation at individual synapses have been intensively studied, whether and how basal release probability variability is collectively controlled across a synapse population is poorly understood, despite its implications in information processing and memory storage (Barbour et al., 2007; Buzsáki and Mizuseki, 2014; Bromer et al., 2018; Rotman and Klyachko, 2013). Our study highlights the novel contribution of astrocyte GluN2C NMDAR signaling in broadening the range of basal release efficacy of Schaffer collateral synapses by effectively maintaining strong synapses stronger and weaker synapses weaker without an overt effect on the mean synaptic strength.

The present study took advantage of paired-pulse response, a short-term plasticity inversely related to presynaptic release probability as a measure of presynaptic efficacy (Dobrunz and Stevens, 1997), which in some experiments was further corroborated by the correlated changes in CV^-2^ (Malinow and Tsien, 1990; Larkman et al., 1992). This study extends the concept of NMDAR-mediated glial control of PPR (Letellier et al 2016) to include populations of synapses within defined hippocampal subregions in the healthy adult mouse brain. Sampling a small number of synapses by a relatively weak stimulation intensity (i.e. mean EPSC amplitude of ∼60 ± 6 pA to Schaffer collateral stimulation in control condition) consistently revealed a component of PPR distribution that was bi-directionally regulated by astrocyte NMDARs. Notably, such modulation could be masked when sampling a large synapse population that yields stable ensemble responses (Lines et al., 2017). Altogether, results from (i) pharmacological experiments using subtype-specific NMDAR antagonists, which allowed monitoring the effects of NMDAR inhibition in the same synapse population, (ii) the genetic interference specifically of astrocyte NMDARs, and (iii) the cellular expression analysis of NMDAR subunits, collectively point to GluN2C NMDAR as the major astrocyte receptor that regulates synaptic strength variability of Schaffer collateral inputs.

### Functional significance of the broad distribution of presynaptic strengths

What might be an advantage in maintaining a highly variable presynaptic efficacy across a synapse population of a given input type? Our mathematical modeling and simulation data indicate that the width of release probability distribution can bias the outcome of activity-dependent synaptic plasticity. In the BCM-like model, a broad release probability distribution favors activity-dependent synaptic depression, while for cases involving a large extent increase in synaptic strength, increased release probability variance promotes potentiation. In a spiking neuron model, the increased release probability variance also promoted LTP while the impact on LTD may depend on the fine structure of the spikes. Notably, several studies have implicated astrocyte signaling in LTD in hippocampal CA3 to CA1 synapses (Chen et al., 2013; Andrade-Talavera et al., 2016; Navarrate et al., 2019; Pinto-Duarte et al., 2019), where spike timing-dependent LTD in slices from young mice is sensitive to inhibitors of GluN2C/D, although the source of GluN2C/D remains to be determined (Andrade-Talavera et al., 2016). Astrocyte GluN2C NMDAR-dependent maintenance of variable basal presynaptic strengths of CA3 to CA1 synapses that we have identified here could therefore be linked to astrocyte signaling that promotes LTP and LTD, which in turn, are not only important for learning and memory but also for network stability (Collingridge et al., 2010; Royer and Paré, 2003; Zenke and Gerstner, 2017). Curiously, this GluN2C NMDAR-dependent broadening of the basal presynaptic efficacy in CA1 neurons is confined to the SR inputs and not observed for SO or SLM inputs. This suggests that for dendritic computations in CA1 pyramidal neurons, the broad variability of synaptic strengths at CA3-CA1 synapses is more crucial compared to the synaptic strength variability of basal or apical tuft inputs. Mice deficient in GluN2C, while mostly normal in their behavior, show deficits in acquisition of conditioned fear and working memory and changes in neuronal oscillations (Hillman et al., 2011; Mao et al., 2020). Some population of interneurons express GluN2C, however (Gupta et al., 2016; Ravikrishnan et al., 2018), and this confounds the interpretation of the observed effects solely to deficits in astrocyte GluN2C. It would be of interest to determine in the future whether there is a learning performance deficit in mice specifically deficient in astrocyte GluN2C or GluN1 in the hippocampal CA1 subfield.

The mechanism by which astrocyte GluN2C NMDARs are coupled to the changes in presynaptic efficacy remains to be clarified. The narrowing of the range of PPR suggest a bi-directional modulation, in one possibility, that astrocyte NMDAR signaling could trigger the release of two types of gliotransmitters with one potentiating and another depressing (Schwarz et al., 2017; Covelo and Araque, 2018). Interestingly, astrocyte-mediated release of ATP, which is converted to adenosine by extracellular ATPases, has been implicated in the bi-directional modulation of presynaptic efficacy (Panatier et al., 2011; Pascual et al., 2005; Zhang et al., 2003; Tan et al., 2017). The effects of adenosine depend in part on the presynaptic A_2A_ or A_1_ receptors that either enhance or suppress presynaptic function, respectively (Panatier et al., 2011; Tan et al., 2017). Similarly, the relative abundance of A_2A_ or A_1_ receptors at individual presynaptic boutons could determine the polarity of presynaptic efficacy upon accumulation of extracellular adenosine by the astrocyte GluN2C NMDAR activity. In another scenario, astrocyte GluN2C receptors may influence the release of other gliotransmitters such as glutamate (Jourdain et al., 2007) to target presynaptic or postsynaptic glutamate receptors and/or GluN2C receptor signaling may influence astrocyte-mediated K^+^ clearance to locally shape presynaptic action potential waveforms (Cui et al., 2018) to in turn affect presynaptic efficacy.

### Astrocyte NMDAR subunit mRNAs and implications for layer-specific synapse modulation

Previous transcriptome analysis of the major cell types in mouse cerebral cortex have suggested that the GluN2C NMDAR subunit mRNA is one of the highly enriched transcripts in astrocytes, whose level can be up to 70 fold of the level found in neurons (Zhang et al., 2014). Our single cell RT-PCR analyses also show robust expression of GluN2C mRNA in astrocytes across the three CA1 layers in contrast to GluN2C mRNA expression in pyramidal neurons. Curiously, despite the high expression of GluN2C mRNA in astrocytes, only low levels of GluN1 mRNA is detected in astrocytes. This finding is unexpected given that GluN1 subunit is required for the surface expression of functional NMDARs (Fukaya et al., 2003; Abe et al., 2004). Moreover, the observed occlusion of QNZ46 effects on synaptic transmission by astrocyte-specific knock-down of GluN1 also support the presence of functional heteromeric GluN1/GluN2C NMDARs in astrocytes. The whole-cell patch clamp method we used to collect RNAs is biased towards sampling of transcripts that are abundant in the cell body. Given that astrocyte processes are numerous and thin, the discordance between the detected GluN1 and GluN2C mRNA levels could be explained if the mRNAs are differentially localized, with GluN1 mRNA being preferentially targeted to processes compared to the cell body of astrocytes. Such a proposal is consistent with a recent study reporting of GluN1 mRNA in astrocyte processes that is locally translated (Sakers et al., 2017). The precise intracellular localization of NMDAR subunit mRNAs in astrocytes remains to be determined.

The present study revealed synapse regulation by the astrocyte GluN2C NMDARs that is confined to the SR input although GluN2C mRNA is expressed broadly across CA1 astrocyte layers. Therefore, the layer-specificity of synaptic modulation could potentially arise from features of GluN2C NMDAR assembly and trafficking and/or signaling that is unique to SR astrocytes over SO and SLM astrocytes. In support of layer-specific differences in astrocyte NMDAR signaling, the slow astrocyte membrane depolarization triggered by puff applied NMDA-glycine shows significant dependence on astrocyte NMDAR only in SR and not in SO nor in SLM (Figure S6). Additionally, differences in the properties of presynaptic inputs to CA1 pyramidal neuron dendrites across layers (e.g. Schroeder et al., 2018) could also contribute to layer-specific synaptic modulation by astrocyte GluN2C NMDARs.

The GluN2C subunit forms diheteromeric receptor complexes with the obligate GluN1 subunit and triheteromeric receptor complexes with GluN1 and GluN2A (Paoletti et al., 2013; Hansen et al., 2018). The presence of GluN2C confers NMDAR properties that are distinct from NMDARs containing GluN2A or GluN2B which are abundant in neurons. For example, GluN2C-containing NMDARs show reduced channel open probability, increased glutamate sensitivity, slow receptor deactivation, and a decreased sensitivity to Mg^2+^ block (Paoletti et al., 2013; Hansen et al., 2018). Weak Mg^2+^ binding would enable open-pore blockers such as MK801, to nonetheless rapidly exert their inhibitory action. Such reduced Mg^2+^ sensitivity of astrocyte NR2C NMDARs could have contributed to the rapid effect observed for MK801 in normalizing presynaptic strengths and reducing the PPR disparity. Moreover, the low EC_50_ of GluN2C-containing NMDAR activation to glutamate (Hansen et al., 2018) suggests that they may be well suited for detecting synaptic release events at perisynaptic astrocyte processes that can be at some distance from the active zone. Several drugs that target NMDARs have been in clinical use, such as ketamine as an anesthetic and treatment for depression (Krystal et al., 2019; Williams and Schatzberg, 2016) and memantine for the treatment of moderate to severe dementia in Alzheimer’s disease (Graham et al., 2017). Under physiological conditions, GluN2C/GluN2D-containing NMDARs display up to 10-fold higher sensitivity to ketamine and memantine in comparison to GluN2A/GluN2B-containing NMDARs that are highly expressed in neurons (Kotermanski and Johnson, 2009; Hansen et al., 2017). Therefore, although therapeutics targeting NMDARs to date have largely focused on neuronal NMDARs, it would be crucial to consider also the consequences of interfering with GluN2C NMDARs that are enriched in astrocytes, which is underscored by the increasing recognition of the involvement of astrocytes in a variety of neurological disorders (Zuchero and Barres, 2015; Chung et al., 2015).

## Materials and Methods

### Slice preparation and whole cell recordings

Mice (P60-120) were deeply anesthetized with isoflurane and transcardially perfused with ice-cold cutting aCSF containing (in mM) 93 N-methyl D-glucamine, 2.5 KCl, 1.2 NaH_2_PO_4_, 30 NaHCO_3_, 20 HEPES, 20 glucose, 5 Na ascorbate, 2 thiourea, 3 sodium pyruvate, 12 N-acetyl L-cysteine, 10 MgSO_4_, 0.5 CaCl_2_, pH adjusted to 7.4 with HCl and bubbled with 95% O_2_ / 5% CO_2_. Brains were extracted and 350 μm transverse hippocampal slices were cut in ice-cold cutting aCSF on a Leica VT1200 vibrating microtome. Slices were incubated for 12 min in cutting aCSF warmed to 34°C, then placed in holding solution containing (in mM) 81.2 NaCl, 2.5 KCl, 1.2 NaH_2_PO_4_, 30 NaHCO_3_, 20 HEPES, 20 D-glucose, 5 Na ascorbate, 2 thiourea, 3 sodium pyruvate, 12 N-acetyl L-cysteine, 2 MgSO_4_, 2 CaCl_2_, pH 7.4, bubbled with 95% O_2_ / 5% CO_2_ for up to 8 h. In some cases, slices were incubated in 50 µM sulphorhodamine for 30 min at 34°C to identify astrocytes for RNA extraction. Whole-cell patch clamp recordings were obtained from CA1 pyramidal neurons or astrocytes using an Olympus BX51W1 microscope equipped with IR-DIC optics and a motorized stage. Images were captured with a digital camera (Andor, iXion3) and imaged using MetaMorph software (Molecular Devices). Voltage clamp and current clamp experiments were carried out using Multiclamp 700B amplifiers (Molecular Devices) and up to four micromanipulators (SM5, Luigs and Neumann). Patch pipettes (tip resistance 2-4 MΩ for neurons, 4-6 MΩ for astrocytes) were pulled using a vertical 2-stage puller (PC-10 Narishige). Slices were constantly perfused with recording aCSF containing (in mM) 119 NaCl, 2.5 KCl, 1.3 NaH_2_PO_4_, 26 NaHCO_3_, 1 MgCl_2_, 2 CaCl_2_, 20 D-glucose and 0.5 Na ascorbate pH 7.4, bubbled with 95% O_2_ / 5% CO2 and maintained at 32-34°C using an in-line heater (Harvard Instruments). Picrotoxin (100 µM) was added to the recording aCSF to block GABA_A_-receptors and isolate glutamatergic synaptic transmission. Internal pipette solution contained (in mM) 130 CsMeSO_3_, 8 NaCl, 4 Mg-ATP, 0.3 Na-GTP, 0.5 EGTA, 10 HEPES, pH 7.3, 290-295 mOsm for neuron recordings, and 130 K gluconate, 10 HEPES, 4 MgCl_2_, 4 Na_2_-ATP, 0.4 Na_3_-GTP, 10 Na-phosphocreatine, pH 7.3, 290 mOsm for astrocyte recordings. 50 µM AlexaFluor 488 or 594 hydrazide (Thermo Fisher Scientific) was included in the pipette solution to verify cell identity and to position stimulating and puff pipettes. Neurons were voltage clamped at −70 mV and series resistance (R_s_) was monitored throughout all recordings and left uncompensated. Experiments were discarded if R_s_ <30 MΩ and/or changed by >20%. Average R_s_ was 25.24 ± 0.27 MΩ (n=298 cells).

### Recordings of synaptic transmission

For all experiments using NMDAR antagonists, 1 mM MK801 was included in the patch pipette solution. Internal solution was allowed to equilibrate into the cell for 10-15 min and inputs were stimulated at a low frequency (0.1Hz) with pairs of pulses at least 45 times before beginning the experiment to pre-block postsynaptic NMDAR receptors. Effective inhibition of NMDAR currents by internal MK801 was confirmed in a separate set of experiments (Figure S1). Pipettes were tip filled with ∼0.2 µl of internal solution lacking MK801 to avoid its leakage to the extracellular milieu prior to seal formation. Small bundles of axons in the *stratum radiatum*, *oriens*, or *lacunosum moleulare* were stimulated using AgCl bipolar electrodes in theta-glass pipettes (tip diameter ∼2-3 µm) filled with recording aCSF, connected to a stimulus isolation unit (A360, WPI). These bundles are identified throughout as ‘input’. Stimulation strengths varied between ∼50-500 µA and were adjusted to obtain EPSCs of approximately 50-100 pA. Up to three independent inputs were sampled during a single experiment, though a maximum of two independent inputs were activated per input pathway. When multiple stimulation electrodes were used they were positioned on opposite sides of the neuron and/or in different input pathways (i.e. SR and SLM, or SR and SO). When sampling two inputs in a single pathway, the independence of the two inputs was confirmed by a cross-paired-pulse stimulation paradigm (Letellier et al., 2016); after stimulating one of the two inputs the other input was stimulated in quick succession (50 ms inter-pulse interval). If facilitation or depression was observed in the second pulse, then the position of stimulation electrode was changed and independence of the two inputs was re-assessed. During the experiment, EPSCs from each pathway were sampled with paired pulses every 30 s. Inputs from separate pathways were stimulated at least 5 s apart. EPSCs were sampled over a 10 min baseline period (i.e. 20 sweeps) before the perfusion of drugs and continued for at least an additional 20 min. EPSC amplitudes were averaged over 20 pre-drug baseline sweeps, and 20 post-drug sweeps. The change in EPSCs (ΔEPSCs) was calculated as the ratio of the average post-drug amplitude to the average baseline amplitude. PPRs were calculated based on the average EPSCs of 5 sweep bins (i.e. 2.5 min). Baseline and post-drug averages were calculated as the average of four bins (i.e. over 10 min) before and after drug application, respectively. Changes in PPR (ΔPPR) were calculated as the difference between the drug application value and the baseline value. Values for coefficient of variation (CV) were obtained from 20 baseline, and 20 post-drug sweeps. The change in CV^-2^ (ΔCV^-2^) was calculated as the ratio of post-drug CV^-2^ to pre-drug CV^-2^. EPSCs and sEPSCs amplitudes in the absence of drug application were stable over the duration of the experiment, and sEPSC amplitudes were stable across all drug conditions tested (Figure S3, S6), suggesting that run-down (i.e. non-stationarities) will not substantially influence the outcome of the CV^-2^ analysis. All EPSC amplitude, rise-time, and decay measurements were performed in Clampfit 10.6 software. sEPSCs were identified as events outside a 50ms window following the second stimulation pulse for each pathway using the template matching algorithm in Clampfit 10.6.

### Astrocyte recordings and NMDA/glycine puff

Astrocytes were identified in acute slices based on mCherry or sulphorhodamine fluorescence, or on their appearance under IR-DIC observation (small, circular cell bodies in the neuropil). Astrocyte identity was always confirmed by their passive electrical properties (linear I-V relationship), low input resistance (<20 MΩ), and low resting membrane potential (< −75 mV), as well as post-hoc labeling by Alexa dyes included in the patch pipette. Recording aCSF contained picrotoxin (100 µM), tetrodotoxin (0.5 µM), and CNQX (10 µM) to reduce network excitability associated with the application of iGluR agonists. Patch pipettes (R_t_= 4-6 MΩ) were used to locally deliver recording aCSF solution containing 1 mM NMDA and 1 mM glycine. Puff pipettes were placed approximately 50 µm from the patched cell and puff pressure (3 psi, 100 ms duration) was controlled with a Picospritzer III (Parker Hannifin) connected to N_2_ gas.

### Whole-cell patch RNA extraction from astrocytes and neurons

Whole-cell patch clamp recordings from astrocytes and neurons were performed using pipettes containing (in mM) 130 K gluconate, 10 HEPES, 4 MgCl_2_, 4 Na_2_-ATP, 0.4 Na_3_-GTP, 10 Na-phosphocreatine, 1U/µL RNAase inhibitor, 50 µM AlexaFluor488, pH 7.3, 290 mOsm. Pipettes were tip filled (∼0.2 µl) with the same internal solution but lacking RNAase inhibitor in order to facilitate obtaining of GΩ seals. After determining the electrical properties of the patched cell to confirm its identity, RNA was extracted using a previously published protocol (Fuzik et al., 2015). The cell was held at −5 mV, and repetitively depolarized to +20 mV for 5 ms at 100 Hz for ∼5 min while light negative pressure was applied to the pipette. The AlexaFluor fluorescence signal was visualized to confirm the extraction of cell cytoplasm.

### Single cell quantitative PCR

Individual patched cells were processed following the provideŕs recommendation for the Single Cell-to-CT™ qRT-PCR Kit (ThermoFisher Scientific). cDNAs for GRIN1, GRIN2A, GRIN2B, GRIN2C, GRIN2D and Rn28s1 were quantified by TAQMAN system using the following probes. The provider’s recommended pre-amplification step was performed for all the genes except for Rn28s1.

GRIN1 fw 5’GACCGCTTCAGTCCCTTTGG 3’, rv 5’ CACCTTCCCCAATGCCAGAG 3’, probe 5’ AGCAGGACGCCCCAGGAAAACCAC 3’ (MGBNFQ, 6FAM)

GRIN2A fw 5’ AGACCCCGCTACACACTCTG 3’, rv 5’TTGCCCACCTTTTCCCATTCC 3’, probe 5’AGCACGATCACCACAAGCCTGGGG 3’ (MGBNFQ, 6FAM)

GRIN2B fw 5’GGCATGATTGGTGAGGTGGTC 3’, rv 5’GGCTCTAAGAAGGCAGAAGGTG 3’, probe 5’ATTGCTGCGTGATACCATGACACTGATGCC 3’ (MGBNFQ, 6FAM)

GRIN2C fw 5’GGAGGCTTTCTACAGGCATCTG 3’, rv 5’ATACTTCATGTACAGGACCCCATG 3’, probe 5’TCCCACCGTCCCACCATCTCCCAG 3’ (MGBNFQ, 6FAM)

GRIN2D fw 5’TCAGCGACCGGAAGTTCCAG 3’, rv 5’TCCCTGCCTTGAGCTGAGTG 3’, probe 5’ TCCTCCACTCTTGGCTGGTTGTATCGCA 3’(MGBNFQ, 6FAM)

Rn28s1 fw 5’CCTACCTACTATCCAGCGAAACC 3’, rev 5’AGCTCAACAGGGTCTTCTTTCC 3’, probe 5’ CTGATTCCGCCAAGCCCGTTCCCT 3’ (MGBNFQ, VIC)

RNA levels were normalized by the quantity of Rn28s1, and standard curves were prepared for estimating the quantity (in femtograms) of the targeted RNAs after pre-amplification. For making the standard curves, total RNA from mouse brain hippocampal tissue was extracted using TRIZOL reagent. Target cDNAs were amplified by regular PCR, and amplicons were purified, quantity of cDNA was measured and submitted to serial dilutions, used for standard curves in TAQMAN system, in parallel to the single cell derived cDNA samples.

### Intracranial AAV injections

Recombinant AAV vectors were targeted to the dorsal CA1 hippocampus of adult (P60-P90) GRIN1^flx/flx^ mice as previously described (Letellier et al., 2016; Cetin et al., 2007). Bilateral injections (∼600 nl/hemisphere at a rate of 200 nl/min) of AAV9.2-GFAP104-nls-mCherry-Cre (6.67⨯10^13^ genome copies/ml), AAV9.2-GFAP104-nls-mCherry (i.e. ΔCre; 6.86⨯10^13^ genome copies/ml), AAVDJ8-GFAP104-nls-mCherry-Cre (5.95⨯10^13^ genome copies/ml), or AAVDJ8-GFAP104-eGFP (also denoted ΔCre for simplicity; 3.98⨯10^13^ genome copies/ml) were made using the following coordinates; X (posterior from Bregma) – 1.4 mm, Y (lateral from sagittal suture) ± 1.4 mm, Z (ventral from pia) – 2.1 mm. Viruses were expressed for at least 21 days (max post-surgical duration of 40 days) before mice were used in physiology or imaging experiments.

### Immunohistochemistry

Slices prepared from GRIN1*^flx/flx^* mice expressing GFAP104-nls-mCherry-Cre were fixed in 4% paraformaldehyde (PFA) in 0.1 M phosphate buffer solution (PB) for 1-2 h, then washed in PB and stored overnight. Alternatively, mice brain tissue was fixed with 4% PFA in PB (pH7.5) by transcardial perfusion, followed by further overnight incubation of the removed brain. Fixed brains were cryosectioned to a thickness of 40 μm. Slices were permeabilized with 0.3% Triton X-100 and blocked in 10% goat serum, then incubated in primary antibody overnight at 4°C. Slices were washed and incubated in secondary antibodies for 1-2 h at room temperature. Slices were then washed and mounted in ProLong antifade mounting medium (ThermoFischer Scientific) containing DAPI (1:1000, ThermoFisher Scientific), and visualized on a Zeiss 780, or an Olympus FV1200 or FV3000 confocal microscope. The primary antibodies used were chicken anti-RFP (1: 1000, Rockland #600-901-379), mouse anti-GFAP (1:1000, Synaptic Systems #173011), rabbit anti-GFAP (1:500, Abcam ab48050), rabbit anti-NeuN (1:500, Abcam #ab177487), and guinea pig anti-GluN1 (1:100, Alamone Labs AGP-046). The secondary antibodies used were AlexaFluor555 goat anti-chicken (1:1000, Invitrogen #A32932), AlexaFluor633 goat anti-mouse (1:1000, Invitrogen #A21052), AlexaFluor 488 goat anti-rabbit (1:1000, Invitrogen #A11034), AlexaFluor 488 goat anti-rabbit (1:500, ThermoFischer Scientific), and AlexaFluor 647 goat anti-guinea pig (1:200, Invitrogen A21450).

### NMDA Receptor immunoprecipitation from mouse brain

Mouse hippocampal tissues were dissociated using a glass dounce homogenizer in a 50 mM Tris-HCl, pH 9.0 buffer containing protease inhibitors (10% w/v, cOmplete^TM^, Merck). Subsequently, 1% (w/v) sodium deoxycholate was added and incubated for 30 min at 37°C with mild shaking to solubilize the tissue. Samples were centrifuged at 100,000 rpm at 4°C for 1h. Supernatant was collected, protein concentration was measured by the BCA assay, and stored at −80°C for later analysis or diluted 5-fold in a 50 mM Tris-HCl pH 7.5 buffer containing 0.1% of Triton X-100 for co-immunoprecipitation experiments.

To test for co-immunoprecipitation, a mix of Sepharose Fast-Flow protein A and protein G beads was prepared. For each antibody reaction, 40 μl of the mixed resin was incubated with 5 μg of antibody. After at least 2 h of incubation at 4°C excess antibody was removed by washing, and 2 mg of protein extract was added to each sample containing the resin beads bound by the antibody and incubated overnight at 4°C. Resins were washed 3 times with 10 volumes of 50 mM Tris-HCl pH 7.5 with 0.1% Triton X-100. Supernatant was carefully removed, and resins were suspended in 2x SDS-PAGE protein loading buffer containing DTT. The unbound protein extract was treated for a second overnight incubation at 4°C with a freshly prepared antibody-bound resin under identical conditions as the first overnight incubation, to ensure effective pull-down of soluble NMDA receptor content from the extract. Co-immunoprecipitated proteins from the two rounds of incubation were pooled together.

Protein samples (inputs and co-immunoprecipitated proteins) were heat denatured for 2 min at 95°C, and subjected to 6% SDS-PAGE separation followed by western blot. Antibodies used were: mouse anti-NR1 (Synaptic Systems #114011), rabbit anti-GluN2A/2B (Synaptic Systems #244003), rabbit anti-GluN2C (generously provided by Dr. Masahiko Watanabe or purchased from Frontier Institute #GLURE3C-RB-AF270), mouse anti-Cre (Merck Millipore clone 2D8) and mouse anti-BirA (Novus biologicals #NBP2-59939).

### Mathematical Model for the Numerical Investigation

We used the facilitation model based on the Tsodyks-Pawelzik-Markram model (Tsodyks et al., 1998) to reproduce ratios of peaks of excitatory postsynaptic current (EPSC) with a 20-Hz spike input. The synaptic dynamics is modeled by [1]:

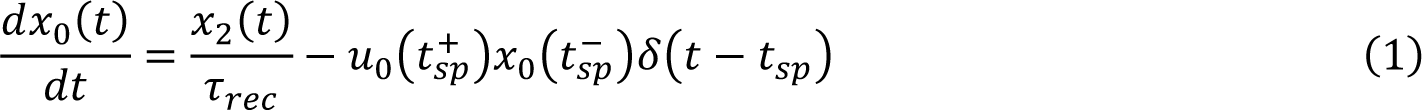

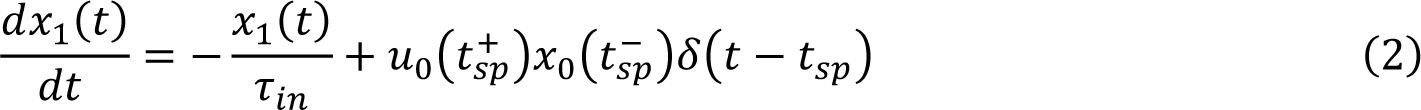

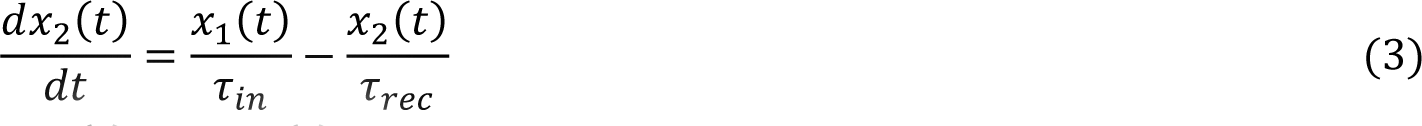

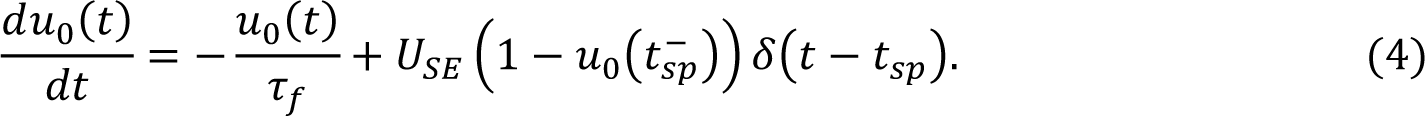

Here x_0_ is the portion of available neurotransmitters, x_1_(t) is the portion of neurotransmitters released by pre-synaptic spikes, x_2_(t) is the portion of neurotransmitters being recovered, u_0_(t) is the utilization of available neurotransmitters after each spike. τ_in_ is the timescale of neurotransmitters release. τ_rec_ is the recovery timescale. τ_f_ is the timescale of synaptic facilitation. U_SE_ is the initial release probability without the influence of synaptic facilitation. In this model, the dynamical variables are x_i_ with x_0_ + x_1_ + x_2_ = 1.

There are four parameters to be determined: τ_f_, τ_rec_, U_SE_ and w(0), where w(0) is the initial connection weight between neurons. For simplicity, w(0) is evenly distributed among 10 synapses (Gal et al., 2017). U_SE_ is a random number drawn from a log-normal distribution, in which the distribution is given by

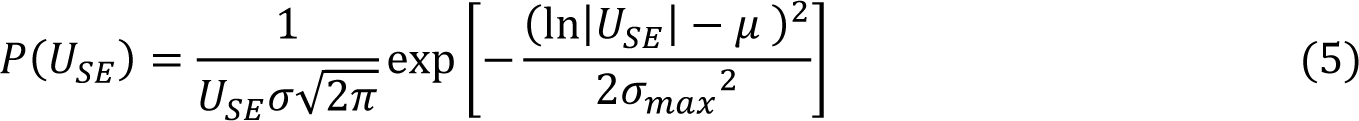

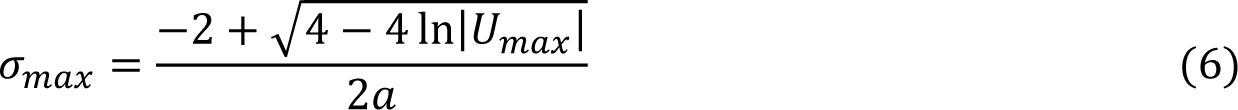

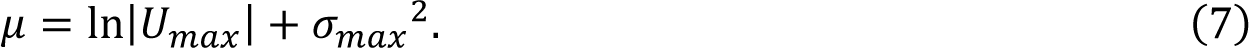

The distribution of release probability in current work was taken from Murthy et al. (1997), which was denoted by U_SE_ . In simulations we use a long-normal function to model the distribution (cf. Figure 3A).

To compare with experimental result, the excitatory postsynaptic current (EPSC) was defined as I(t) = w(0)x_1_(t). The best fit to the experimental data is shown in Figure S8A where the best-fit parameters are τ_f_ = 540 ms, τ_rec_ = 20 ms, U_max_= 0.1 and w(0) = 332.

The BCM-like learning rule for investigation is given by

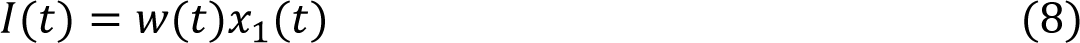

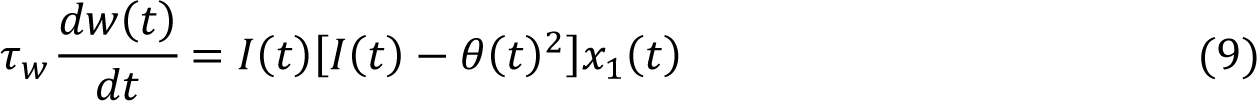

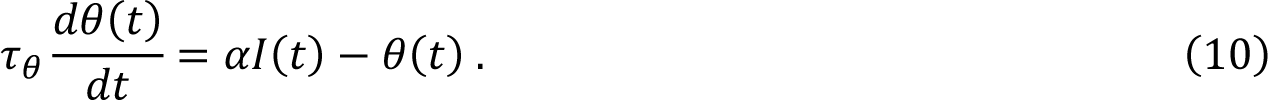

Here θ is the threshold to stop the change of *w*, while τ_θ_ is the timescale to integrate the postsynaptic current I(t). *τ_w_* is the timescale of *w(t)*. α is a parameter to be chosen for different scenarios: LTP or LTD. In this study, the following values were used: τ_w_ = 100 ms and τ_θ_ = 10 ms.

The equations (8) to (10) constitute the BCM learning rule (Bienenstock et al., 1982).

In the series of comparisons shown in Figure 3C, the changes in synaptic plasticity as defined by

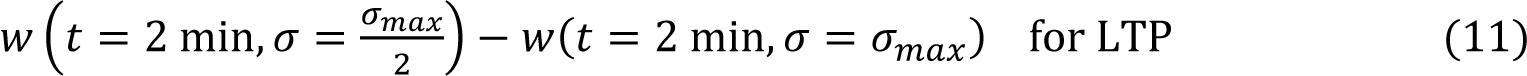

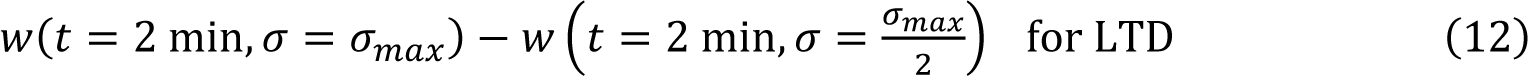

are represented as a function of the absolute change in the connection weight in the control release probability distribution condition as defined by *(w(t = 2 min, σ = σ_max_) − w(t = 0))*. Note that the comparison at t = 2 min was to compare *w(t)* after the epoch of paired stimulations.

Note also that 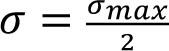
 represents the cases of NMDARs blocked, while σ = σ_max_ represents the control cases. In Figure 3C, the reduction in *σ* always degrades the efficacy of LTD; however, the efficacy of LTP may vary for different level of LTP. For the relative change in *w(t)* ≲ 0.06, the reduction of *σ* does not degrade the LTP efficacy whereas for the relative change in *w(t)* ≳ 0.06, the reduction in *σ* degrades the LTP efficacy.

In the model and simulation based on leaky integrate-and-fire (LIF) neurons, two neurons are connected by 10 synapses. The neurons have time constant ττ = 15 ms, threshold potential V_tre_ = −50.0 mV, reversal potential E_L_ = −65.0 mV and refractory time τ_ref_ = 2 ms. The rising synaptic time constant is 2 ms and decay time constant is ∼5 ms. The synaptic model is the same as equations (1) – (4), and the probability distribution was the same as equations (5) – (7). The best-fit of the model to the data is shown in Figure S8B, where the parameters are τ_f_ = 980 ms, τ_rec_ = 25 ms, U_max_ = 0.1 and w(0) = 6.7.

### Statistics

All statistical analyses were performed using OriginPro software (OriginLab Corp.). Datasets were tested for normality using the Shapiro Wilk test. When the criteria for normality was achieved, differences of mean values were examined using paired or unpaired Student’s two-tailed t-tests or one-way ANOVAs. When the criteria for normality was not achieved, differences in mean values were examined using Mann-Whitney or Kruskal Wallis tests. Normalized mean values obtained from recordings (i.e. normalized EPSCs) were compared to values obtained at the same timepoint in control experiments using Mann-Whitney tests. Variance of PPR distributions were examined using one-tailed f-tests for equal variances or Levene’s test. Box plots represent median and quartile values, whiskers represent maximum and minimum values that are not outliers. P values are indicated throughout or indicated by * if p<0.05.

## Acknowledgements

We would like to thank all members of the Goda laboratory for providing valuable and ongoing feedback throughout the course of this study. We thank Tatjana Tchumatchenko, Thomas Chater and Toru Shinoe for comments on the manuscript at various stages, Yun Kyung Park for help with RNA collection for RT-PCR experiments, Toru Shinoe for pilot electrophysiology experiments, Tom McHugh and Shigeyoshi Itohara for kindly providing *Grin1* floxed mice and Ai9 mice, respectively, and Masahiko Watanabe for generously providing GluN2C antibodies. PC was an Overseas Research Fellow of the Japan Society for the Promotion of Science (JSPS). This work was supported by the RIKEN Center for Brain Science, the Uehara Memorial Foundation, JSPS Core-to-Core Program (JPJSCCA20170008), Grants-in-Aid for Scientific Research (15H04280, YG; 18H05213, TF) from the MEXT, and the Brain/MINDS from the Japan AMED.

## Author Contributions

PHC and YG designed the project and wrote the manuscript. PHC performed electrophysiology experiments and analyzed data. CCAF and TF contributed mathematical modeling and simulation and wrote the manuscript. PHC and AK isolated astrocyte and neuronal RNA for RT-PCR experiments. APF performed biochemistry and RT-PCR experiments. MA and KS made the GluN2C-Cre mice. APF, and AT and MK performed immunohistochemistry experiments. SGG constructed and MK prepared viral reagents.

## Supplementary Figures

**Supplementary Figure S1.**
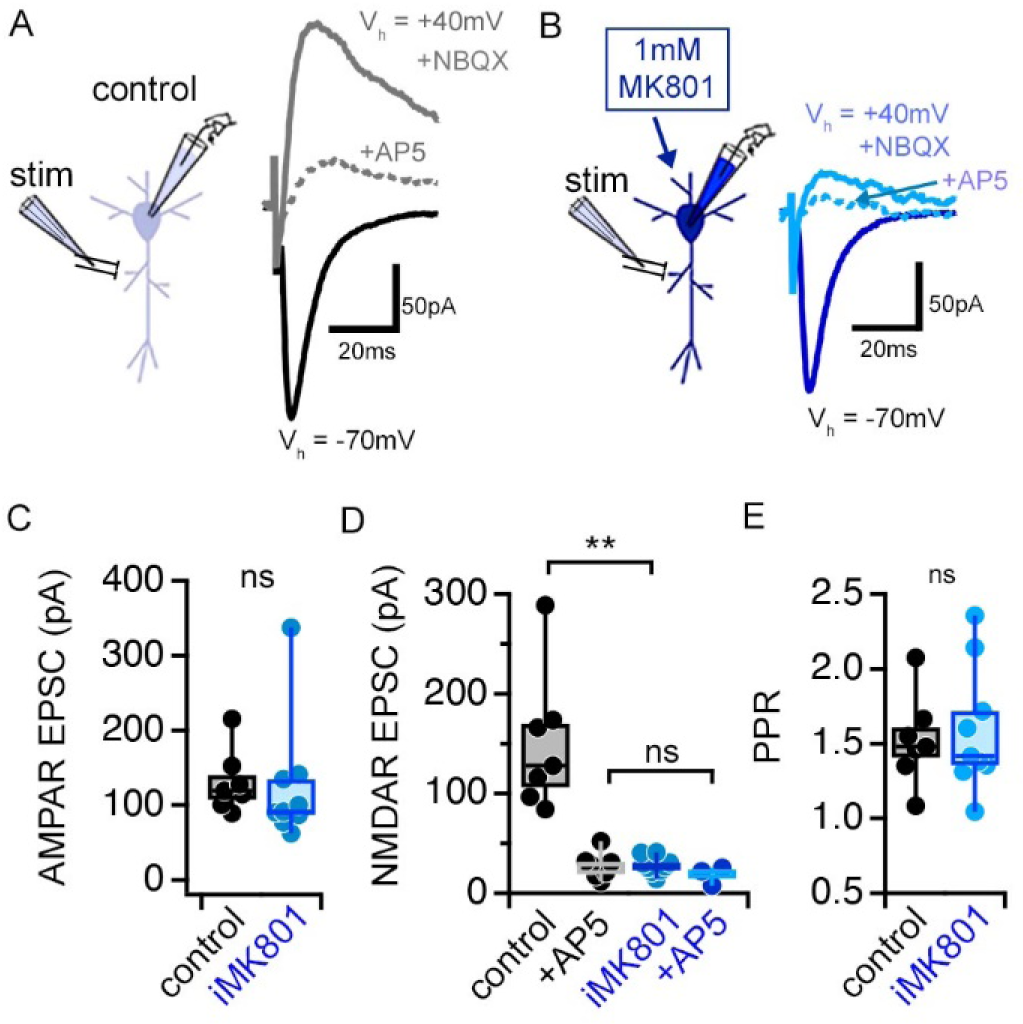
MK801 (1mM) in the patch pipette (iMK801) completely blocks postsynaptic NMDARs. (**A-B**) Representative traces of EPSC recorded at a holding potential (V_h_) of −70mV using control (*A*) or MK801-containing (*B*) pipette solution to measure AMPAR-mediated currents. EPSCs were recorded at a V_h_ of +40mV and in the presence of NBQX (10 μM) to measure isolated NMDAR-mediated currents. AP5 (20 μM) was subsequently added to the bath to confirm the NMDAR-dependence of EPSCs. EPSCs were sampled following a pre-experiment stimulation period that consisted of at least 45 pairs of pulses delivered at 0.1Hz at a V_h_ of −70mV. (**C**) AMPA-mediated EPSCs are no different between control and iMK801experiments. (**D**) NMDAR-mediated EPSCs are completely abolished by the presence of iMK801. (**E**) Paired pulse ratios (PPRs) measured at V_h_ −70mV are not altered by iMK801. Control n = 7 cells, 7 slices, 2 mice; Control +AP5 n = 7 cells, 7 slices, 2 mice; iMK801 n = 9 cells, 9 slices, 3 mice; iMK801 +AP5 n = 3 cells, 3 slices, 2 mice. ns p > .05, ** p < .0001 two-tailed t-test (*D, E*) or Mann-Whitney U test (*C*).

**Supplementary Figure S2.**
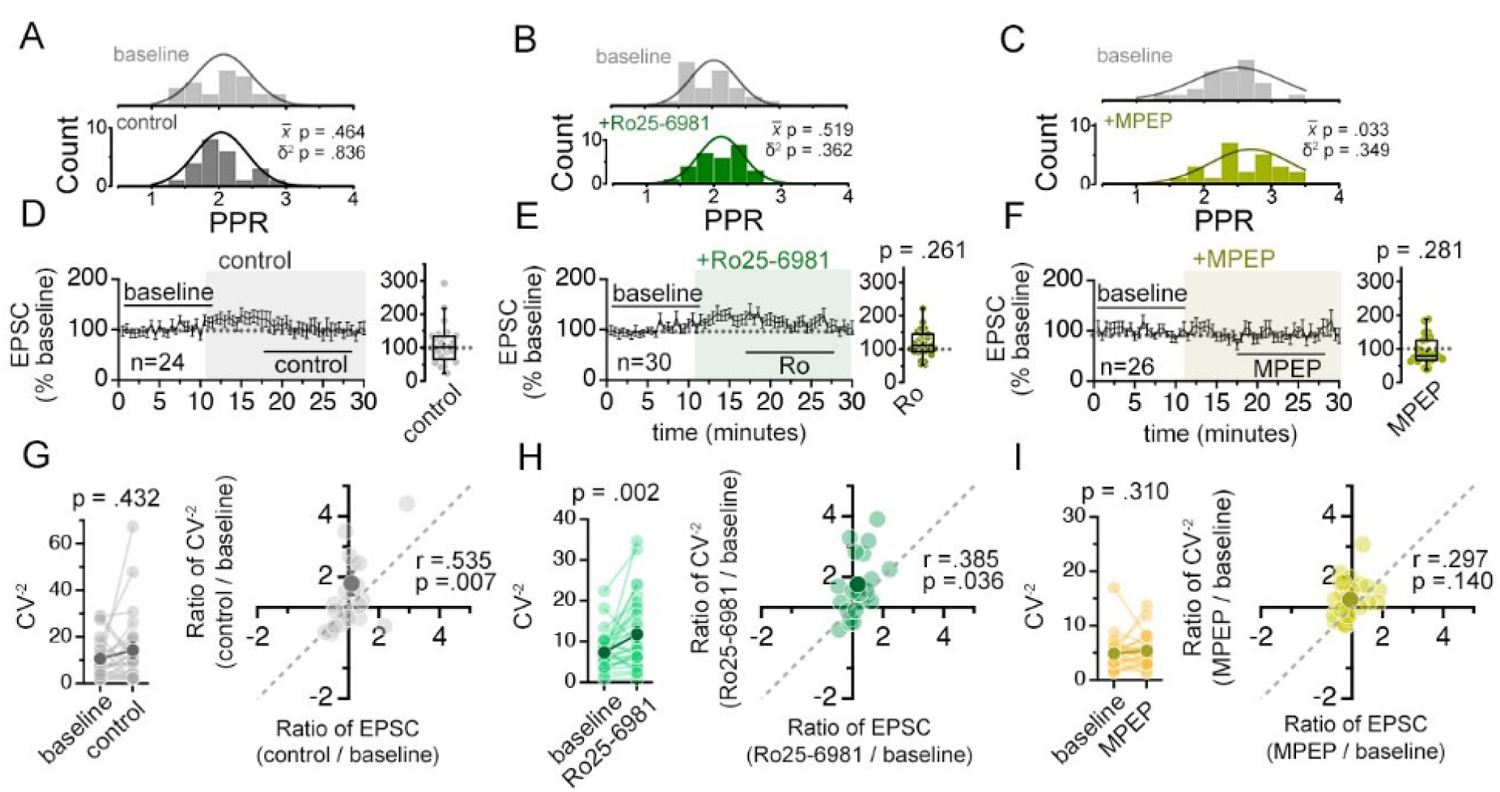
Antagonists of GluN2B-containing NMDARs (Ro25-6981) or mGluR5 (MPEP) do not alter synaptic weight diversity. (**A-C**) Histograms of PPR distributions during the baseline (*top histogram*) and control aCSF or drug application (*bottom histogram*) were fit with single Gaussian curves. The p values obtained for the population mean (x̅) (two tailed paired sample t-test for *A, B*, Wilcoxon signed rank test for *C*) and variance (δ^2^) (one-tailed f-test for equal variances) comparing the baseline to the aCSF/drug application periods are shown in the graphs. (**D-F**) Plots of EPSC amplitudes (normalized to the baseline average) before and during the application of NMDAR antagonists (shaded area) where n is the number of inputs examined. Baseline and experimental periods are indicated (black bars). Box plots to the right show the summary of normalized EPSC amplitudes during the application of indicated inhibitors. P-values are the result of Mann-Whitney U tests comparing EPSC change drug application period to the corresponding period in control experiments. (**G-I**) Left: Mean (dark circles) and individual (light circles) CV^-2^ of EPSCs during the baseline and aCSF/drug application periods. P-values shown are the result of Wilcoxon signed rank tests. Right: Scatterplot of the relative change in CV^-2^ (CV^-2^ _experimental_/CV^-2^_baseline_) against the relative change in EPSC amplitude (EPSC _experimental_/EPSC_baseline_) for the mean (dark circles) and individual inputs (light grey circles). Pearson’s correlation coefficients (r) and p values are shown in the graph. Control n = 12 inputs, 24 cells, 11 mice; Ro25-6981 n = 30 inputs, 17 cells, 11 mice; MPEP n = 26 inputs, 13 cells, 5 mice.

**Supplementary Figure S3.**
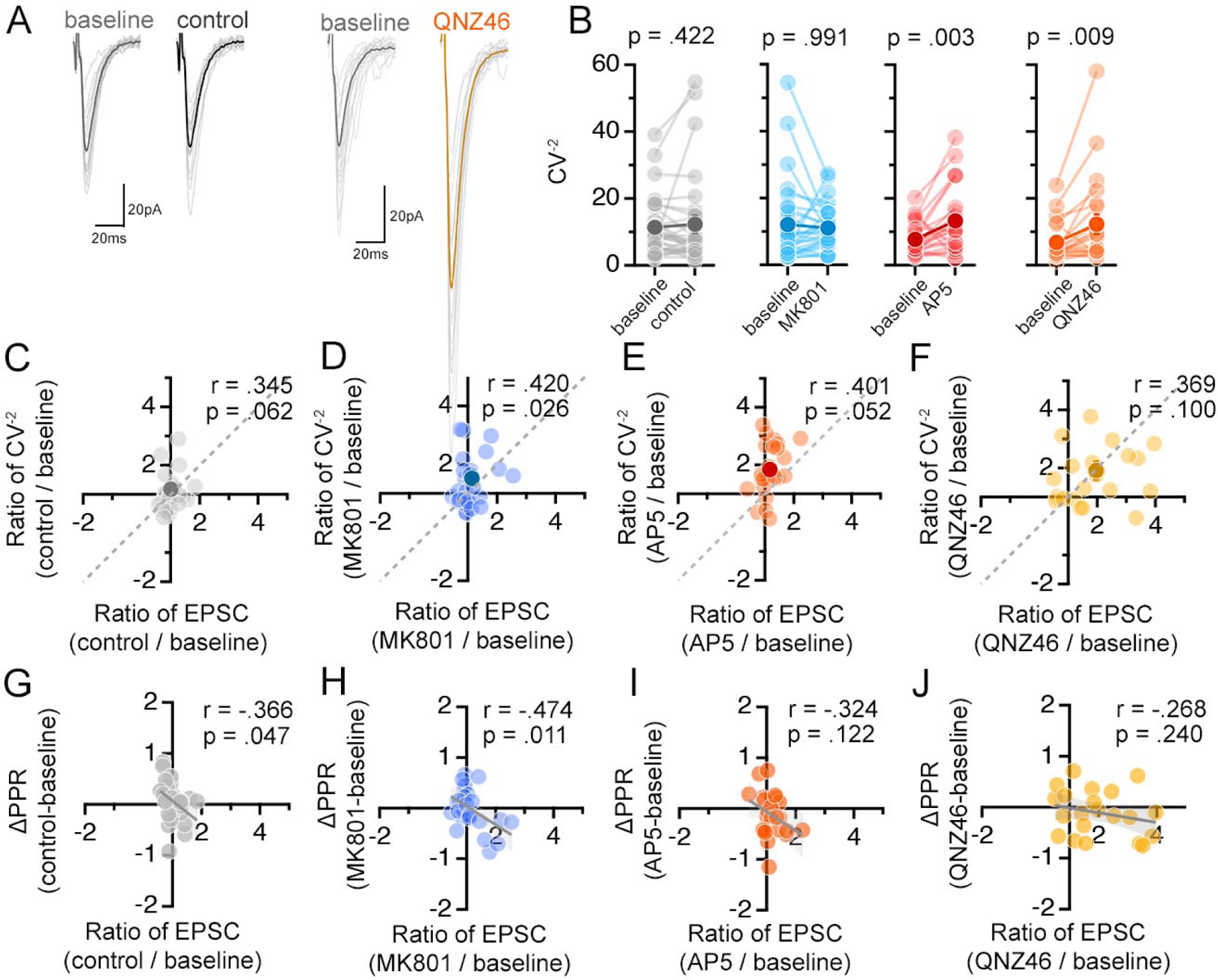
NMDAR antagonists differentially influence presynaptic function. (**A**) Averaged (dark colors) and individual (light grey) EPSC traces during the baseline and experimental periods from a representative experiment. (**B**) Mean (dark circles) and individual (light grey circles) CV^-2^ of EPSCs during the baseline and aCSF/drug application periods. P-values are the result of Wilcoxon signed rank tests. (**C-F**) Scatterplot of the relative change in CV^-2^ (CV^-2^ _experimental_/CV^-2^_baseline_) against the relative change in EPSC amplitude (EPSC _experimental_/EPSC_baseline_) for the mean (dark circles) and individual inputs (light circles). Pearson’s correlation coefficients (r), p values, and the unity line (dashed) are shown in scatter plots. (**G-J**) Scatterplot of the difference in PPR (PPR _experimental_-PPR_baseline_) against the relative change in EPSC amplitude (EPSC _experimental_/EPSC_baseline_) for the mean (dark symbol) and individual inputs (light symbols). Linear fits (grey lines), Pearson’s correlation coefficients (r), and p values are shown in scatter plots. Control = 30 inputs, 15 cells, 13 mice; MK801 = 28 inputs, 14 cells, 9 mice; AP5 = 24 inputs, 12 cells, 11 mice; QNZ46 = 22 inputs, 11 cells, 7 mice; Ro25-6981 n = 30 inputs, 17 cells, 11 mice; MPEP n = 26 inputs, 13 cells, 5 mice.

**Supplementary Figure S4.**
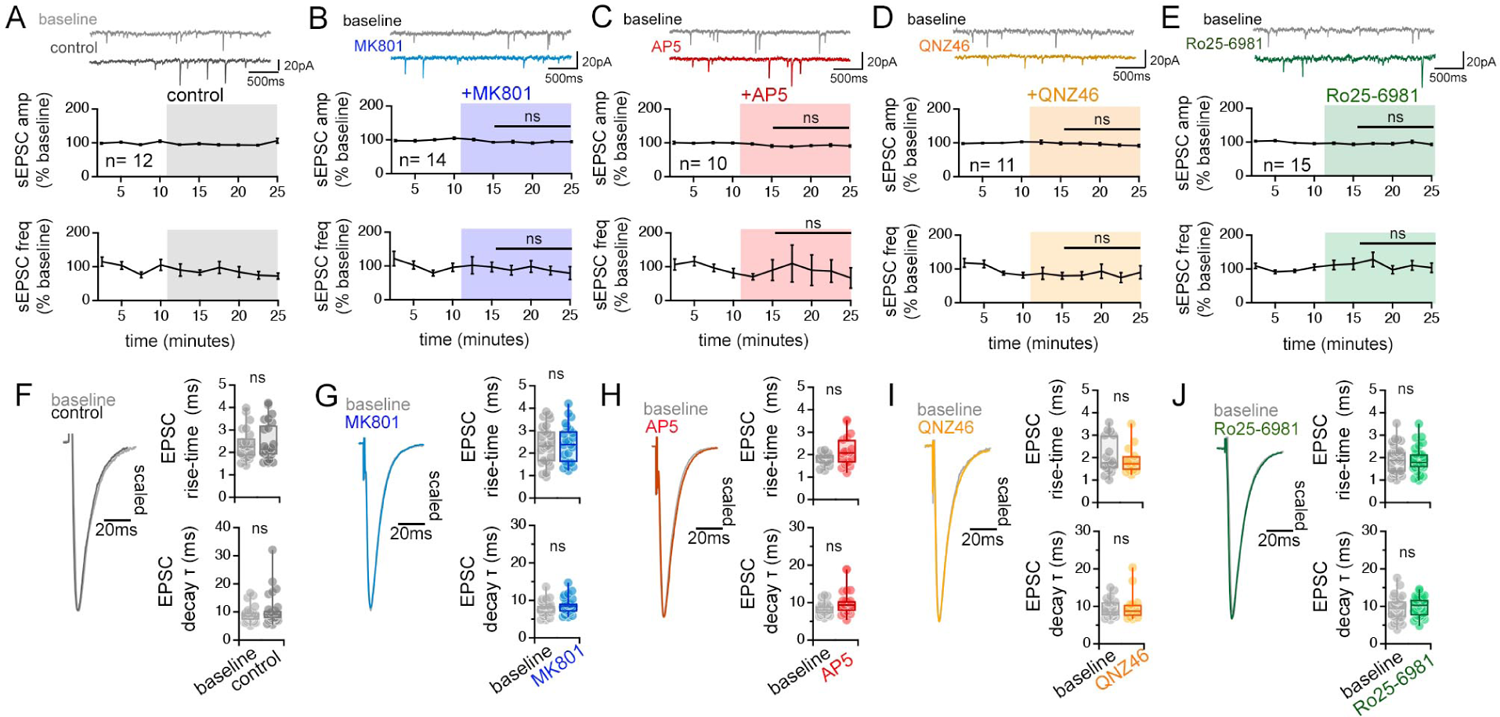
NMDAR antagonists have little effect on spontaneous neurotransmission and the waveform of evoked EPSCs. (**A-E**) Representative traces and quantification of spontaneous EPSC amplitude and frequency recorded during the baseline and after applying control aCSF (*A*), MK801 (*B*), AP5 (*C*), QNZ46 (*D*), and Ro-256981 (*E*). Control n=12 cells, 10 mice; MK801 n = 14 cells, 9 mice; AP5 n = 10 cells, 9 mice; QNZ46 n = 11 cells, 8 mice; Ro25-6981 n = 15 cells, 10 mice. ns p > .05 one way ANOVA (amplitude), and Kruskal-Wallis ANOVA (frequency) comparing the normalized change during experimental periods across treatment conditions. (**F-J**) Left: Grand average waveforms of EPSCs recorded. Right: Box plots of EPSC risetime (top) and EPSC decay time constant (single exponential fit)(bottom) for baseline and control aCSF or drug application. ns p > .05 two tailed t-test or Mann-Whitney U tests. Control n = 24 inputs, 12 cells, 10 mice; MK801 n = 16 inputs, 11 cells, 11 mice; AP5 n = 26 inputs, 13 cells, 9 mice; QNZ46 n = 18 inputs, 9 cells, 6 mice; Ro25-6981 n = 32 inputs, 16 cells, 11 mice.

**Supplementary Figure S5.**
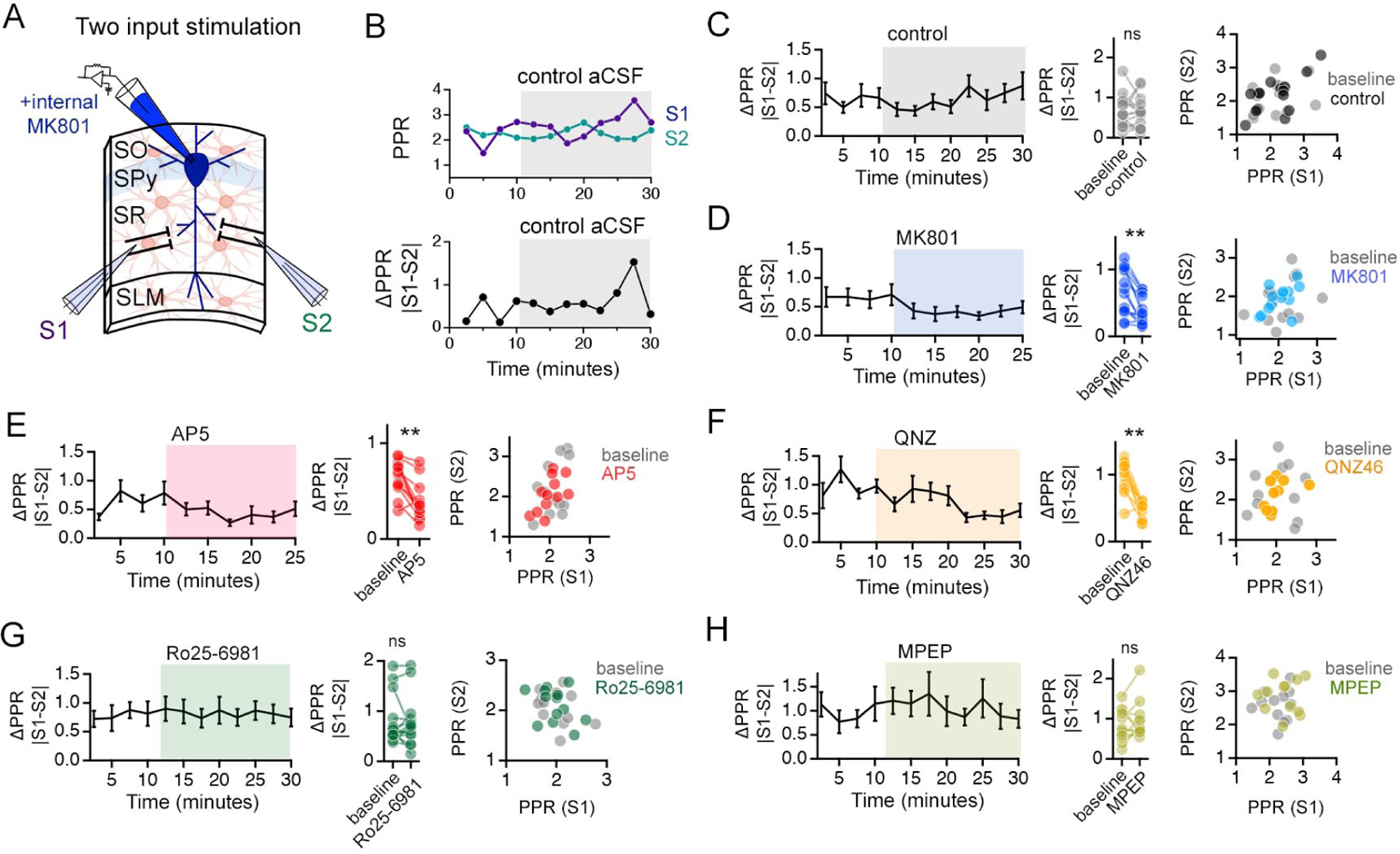
NMDAR antagonists that target GluN2C-containing NMDARs reduce PPR disparity in the *stratum radiatum* (SR). (**A**) Illustration of the two-input stimulation configuration. Two independent inputs in the SR were sampled with interleaved stimulation. (**B**) The absolute difference between the PPRs of the two inputs sampled during a 2.5 min window (average of 5 sweeps) is referred to as ΔPPR (|PPR_S1_-PPR_S2_|) and is taken as a measure of variability of presynaptic strength. (**C-H**) Left: Time course of ΔPPR before and after application of antagonists. Middle: Summary plots of individual experiments comparing ΔPPR in baseline and during antagonist treatment. Right: Scatter plots comparing PPR of the two independent inputs normalized to the first EPSC amplitude for baseline and in the presence of the antagonists. ΔPPR is reduced by MK801 (*D*), AP5 (*E*), QNZ46 (*F*), but not by Ro25-6981 (*G*) or MPEP (*H*). Similarly, the scatterplots demonstrate distribution of average PPR values across pairs of inputs (S1 and S2) of PPRs by MK801, AP5 and QNZ46, all of which target GluN2C-containing NMDARs, but not by Ro25-6981 or MPEP that target GluN2B-NMDAR or mGluR5, respectively. ns p > .05, ** p < .01 Wilcoxon signed ranks test comparing baseline to drug application period. Control n = 15 pairs of inputs, 15 cells, 13 mice; MK801 n = 14 pairs of inputs, 14 cells, 9 mice; AP5 n = 12 pairs of inputs, 12 cells, 11 mice; QNZ46 n = 11 pairs of inputs, 11 cells, 7 mice; Ro25-6981 n = 15 pairs of inputs, 15 cells, 11 mice; MPEP n = 13 pairs of inputs, 13 cells, 5 mice.

**Supplementary Figure S6.**
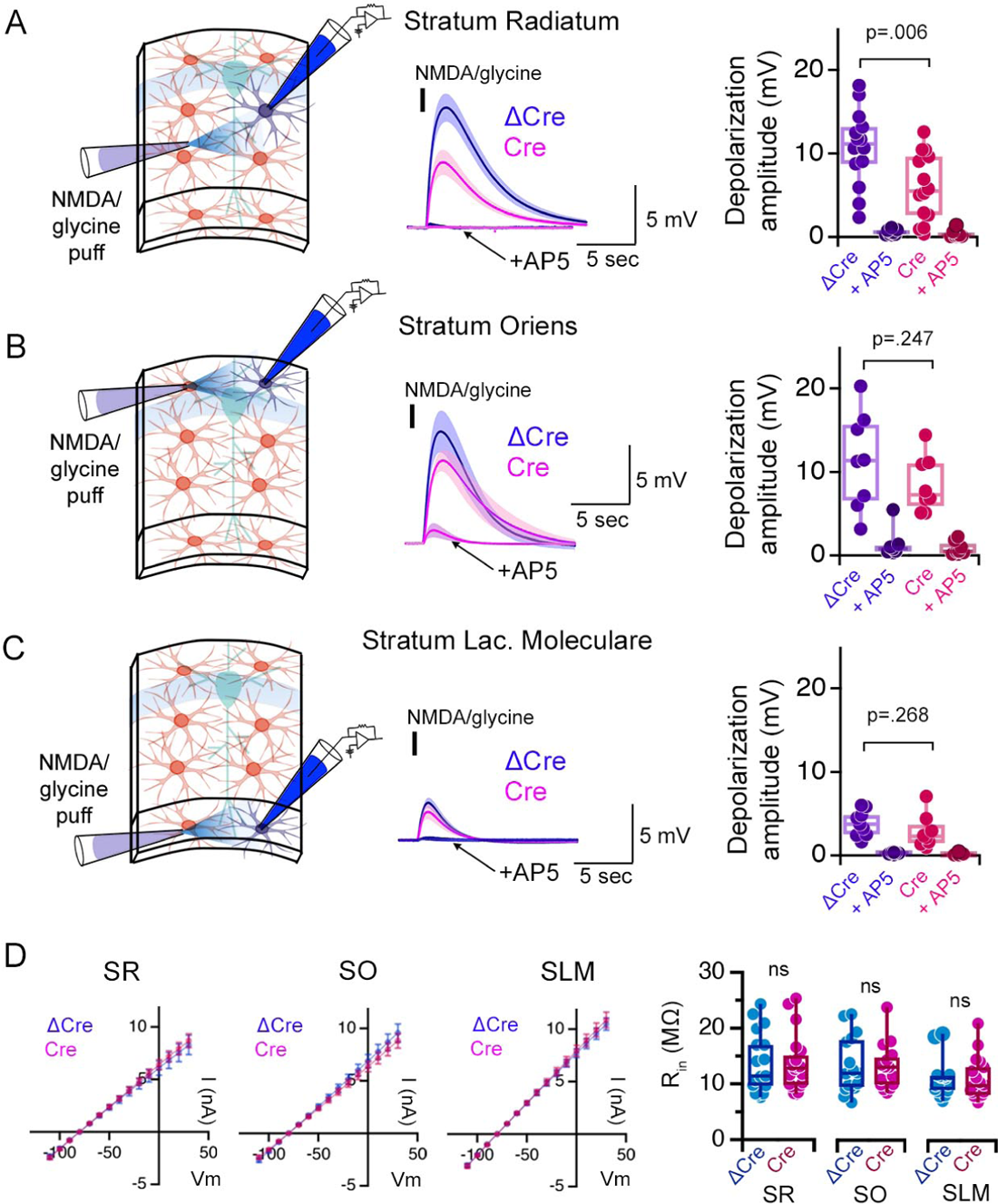
Puff application of NMDA and glycine triggers the depolarization of astrocytes in hippocampal CA1 in acute slices. (**A**) Left: experimental setup. Middle: Local puff application (100 ms, black bar) of NMDA and glycine (1mM each) to astrocytes that are whole-cell patch clamped in the *stratum radiatum* (SR) elicits slow membrane depolarization (ΔCre, blue), which is prevented in the presence of AP5 (arrow). Once the depolarization is triggered, not only NMDARs but also other voltage-gated channels such as L-type calcium channels contribute to the slow depolarization (Letellier et al., 2016). The depolarization is substantially reduced when the same experiment is carried from Cre-expressing astrocytes as monitored by mCherry expression in slices infected with the AAV-GFAP104-nls-mCherry-Cre virus to knock-down astrocyte NMDARs in GRIN1 floxed mice (Cre, red). Right: Summary plot of the peak membrane depolarization in control virus infected (ΔCre) or NMDAR knock-down (Cre) slices in control aCSF or the presence of AP5. (**B-C**) Same as (*A*) except NMDA and glycine puff was applied to astrocytes patch-clamped in the *stratum oriens* (SO)(*B*) and *stratum lacunosum moleculare* (SLM)(*C*). NMDA/glycine puff-mediated depolarizations elicited in SO and SLM astrocytes are not sensitive to astrocyte NMDAR knockout. SR ΔCre n = 14 cells, 3 mice; SR Cre n = 14 cells, 3 mice. SO ΔCre n = 8 cells, 3 mice; SO Cre n = 8 cells, 3 mice. SLM ΔCre n = 9 cells, 2 mice; SLM Cre n = 8 cells, 2 mice. (**D**) Cre expression does not alter the input resistance (R_in_) of astrocytes in any layer. IV relationships of ΔCre and Cre positive astrocytes in the SR, SO, and SLM to square voltage steps (left; 500ms duration) and quantification of R_in_ (right). SR ΔCre n = 17 cells, 8 mice; SR Cre n = 20 cells, 10 mice. SO ΔCre n = 16 cells, 10 mice; SO Cre n = 16 cells, 11 mice. SLM ΔCre n = 19 cells, 8 mice; SLM Cre n = 18 cells, 9 mice. ns p > .05, P values shown are the result of two-tailed t-tests or Mann-Whitney U tests.

**Supplementary Figure S7.**
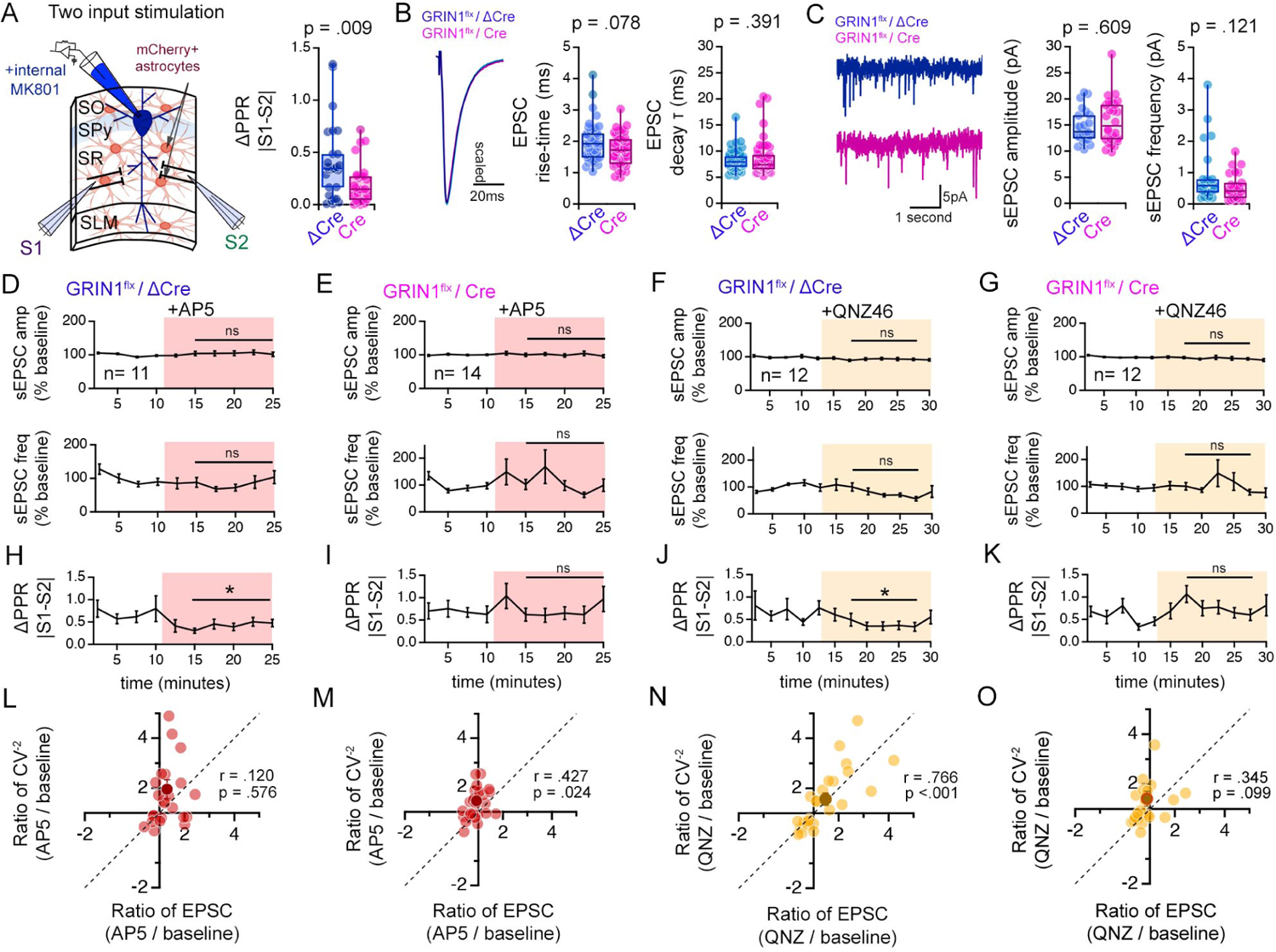
Knock-down of GluN1 in hippocampal astrocytes selectively alters the PPR disparity without affecting the basic properties of synaptic transmission. (**A**) Left: Illustration of the experimental configuration of two-pathway stimulation approach to estimate PPR disparity in slices from GRIN1*^flx/flx^* mice infected with ΔCre or Cre virus. Right: Comparison of ΔPPR (|PPR_S1_-PPR_S2_|) between control ΔCre or Cre slices. PPR disparity (ΔPPR) is reduced in Cre infected slices compared to controls. p = .009, Mann-Whitney U test. ΔCre n = 29 pairs of inputs, 29 cells, 8 mice. Cre n = 28 pairs of inputs, 28 cells, 6 mice. (**B**) Left: Grand average waveforms of EPSCs recorded. Right: Box plots of EPSC risetime and EPSC decay time constant (single exponential fit). Each data point represents the average waveform of at least 5 sweeps of EPSCs. ΔCre n = 46 inputs, 23 cells, 8 mice. Cre n = 41 inputs, 21 cells, 6 mice. (**C**) Left: Representative traces of spontaneous EPSC (sEPSC) recordings from control ΔCre (blue) or Cre (red) slices. Right: Boxplots of sEPSC amplitude and frequency measured in ΔCre or Cre-infected slices. P values shown are the result of Mann-Whitney U test. ΔCre n = 23 cells, 5 mice. Cre n = 26 cells, 5 mice. (**D-G**) Normalized sEPSC amplitude (top) and frequency (bottom) during baseline and in the presence of AP5 or QNZ46 in control ΔCre or Cre-infected slices. Neither sEPSC amplitude nor frequency is not affected by AP5 or QNZ46. Two tailed t-test or Mann-Whitney U test, comparisons made between ΔCre vs. Cre normalized values in post drug period indicated. (**H-K**) ΔPPR (|PPR_S1_-PPR_S2_|) measurements during baseline and in the presence of AP5 or QNZ46 in control ΔCre or Cre-infected slices. The decrease in PPR disparity triggered by AP5 (*H*) and QNZ46 (*J*) is prevented upon knock-down of astrocyte NMDARs. * p < .05 Wilcoxon signed ranks test. ΔCre + AP5 n = 12 pairs of inputs, 12 cells, 6 mice. Cre + AP5 n = 14 pairs of inputs, 14 cells, 5 mice. ΔCre + QNZ46 n = 12 pairs of inputs, 12 cells, 5 mice. Cre + QNZ46 n = 12 pairs of inputs, 12 cells, 6 mice. (**L-O**) Scatterplot of the relative change of CV^-2^ (CV^-2^ _experimental_/CV^-2^_baseline_) against the relative change in EPSC amplitude (EPSC _experimental_/EPSC_baseline_) for the mean (dark symbol) and individual inputs (light symbols). Dashed lines show positive unity line. Pearson’s correlation coefficients (r) and p values are shown in scatter plots. N values are the same as *H-K*. (Supplementary Figure S7 legend continued from previous page)

**Supplementary Figure S8.**
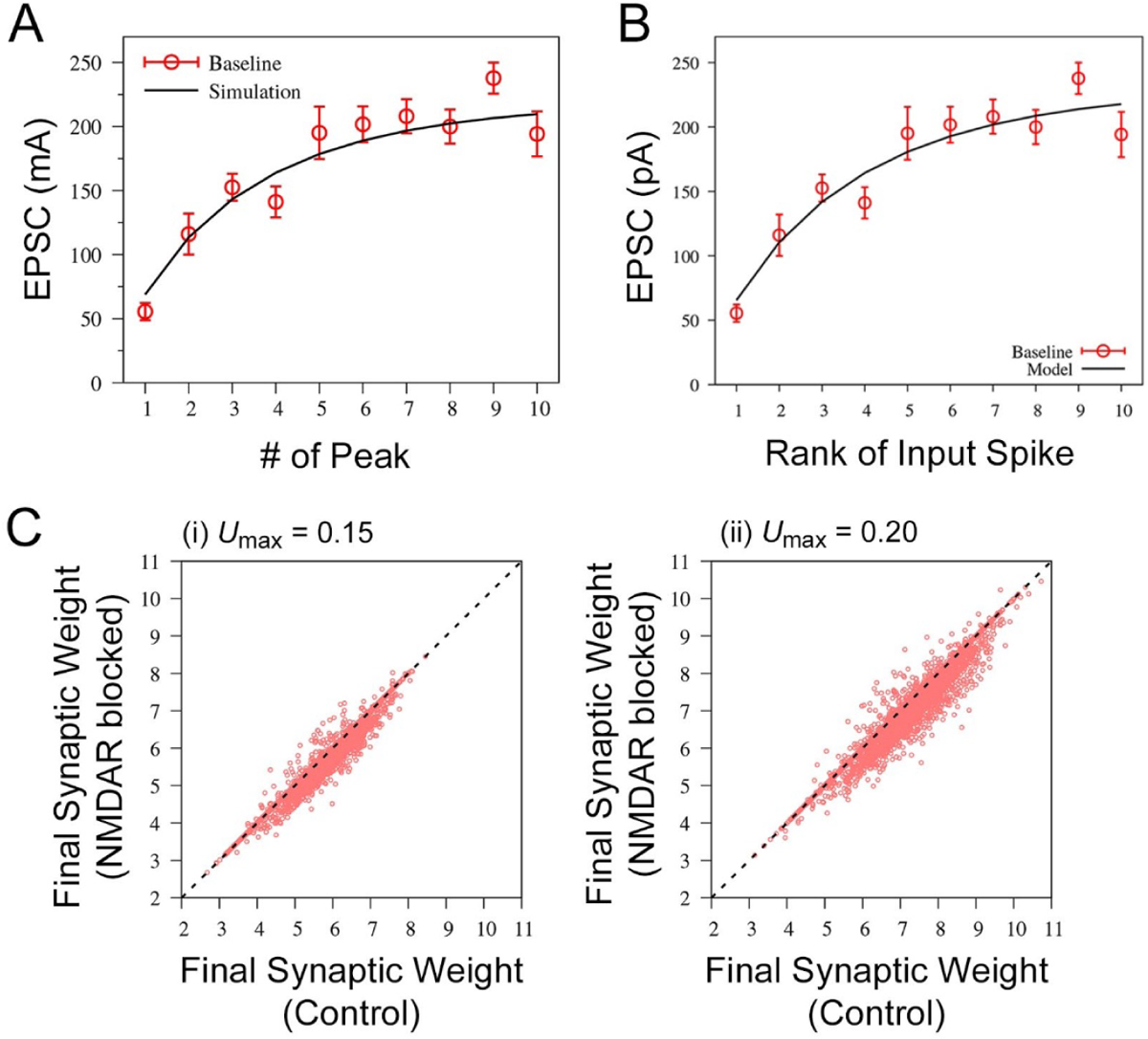
Additional data for mathematical modelling experiments exploring the impact of presynaptic release probability variance on synaptic plasticity. (**A,B**) The best-fits to the experimentally obtained peak EPSC amplitudes of responses triggered by a 20Hz spike train in the BCM-like model (A) and the spiking neuron model (B). The best-fit parameters are τ_ff_ = 540 ms, τ_rec_ = 20 ms, U_max_ = 0.1 and w(0) = 332 in (A), and τ_ff_ = 980 ms, τ_rec_ = 25 ms, U_max_ = 0.1 and w(0) = 6.7 in (B). (**C**) A comparison of the synaptic weight changes observed in response to Poisson spike trains in control condition versus the NMDAR-blocked condition (reduced presynaptic release probability variance) for simulations at (i) *U*_max_ = 0.15 and (ii) *U*_max_ = 0.20. Final synaptic weight above the initial synaptic weight value of w(0) = 6.7, is considered potentiation and below w(0) = 6.7 is considered depression.

**Supplementary Figure S9.**
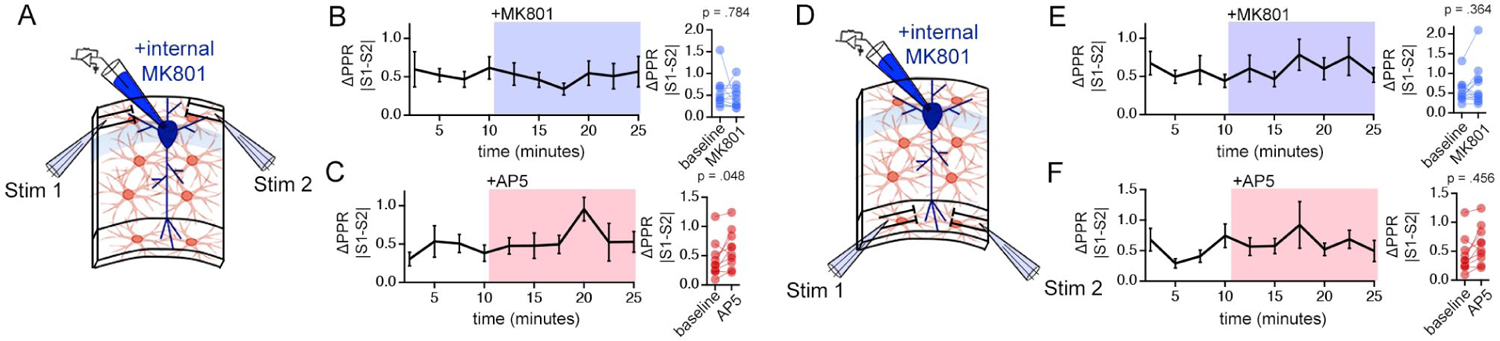
MK801 and AP5 do not alter PPR disparity in the *stratum oriens* (SO) or the *stratum lacunosum moleculare* (SLM). (**A**) Illustration of the two-input stimulation configuration in SO. (**B,C**) Left: PPR disparity [ΔPPR (|PPR_S1_-PPR_S2_|)] during baseline and in the presence of MK801 (*B*) or AP5 (*C*). Right: Summary plot of ΔPPR. Neither MK801 nor AP5 application influences the PPR disparity. (**D-F**) Same experiments as described for (*A-C*) carried out in SLM. Similarly to SO recordings, MK801 and AP5 have no effects of PPR disparity in SLM. P-values are the result of Wilcoxon signed rank tests. SO MK801 n = 12 pairs of inputs, 12 cells, 6 mice; SO AP5 n = 10 pairs of inputs, 10 cells, 7 mice; SLM MK801 n = 13 pairs of inputs, 13 cells, 8 mice; SLM AP5 n = 12 pairs of inputs, 12 cells, 6 mice.

